# Reversible expansion of tissue macrophages in response to macrophage colony-stimulating factor (CSF1) transforms systemic metabolism to fuel liver growth

**DOI:** 10.1101/2023.05.17.538022

**Authors:** Sahar Keshvari, Jesse J.R. Masson, Michelle Ferrari-Cestari, Liviu-Gabriel Bodea, Fathima Nooru-Mohamed, Brian W.C. Tse, Kamil A. Sokolowski, Lena Batoon, Omkar L. Patkar, Mitchell A. Sullivan, Hilmar Ebersbach, Cian Stutz, Robert G. Parton, Kim M. Summers, Allison R. Pettit, David A. Hume, Katharine M. Irvine

**Affiliations:** Mater Research Institute-The University of Queensland, Translational Research Institute, Brisbane, Queensland, 4102, Australia; Clem Jones Centre for Ageing and Dementia Research, Queensland Brain Institute, The University of Queensland, Brisbane, Queensland, 4072, Australia; Preclinical Imaging Facility, Translational Research Institute, Brisbane, Queensland, 4102, Australia; Novartis Institutes for Biomedical Research (NIBR), Fabrikstrasse 2, Novartis Campus, CH-4056 Basel, Switzerland; Institute for Molecular Bioscience, The University of Queensland, Brisbane, Queensland, 4072, Australia; Centre for Microscopy and Microanalysis, The University of Queensland, Brisbane, Queensland, 4072, Australia

**Author notes:** Equal contribution (David A. Hume, Katharine M. Irvine).

**Keywords:** Tissue resident macrophages, innate immunity, metabolic homeostasis, adipose lipid storage, liver, fasting

## Abstract

**Background and Aim:** Macrophages regulate metabolic homeostasis in health and disease. Macrophage colony-stimulating factor (CSF1)-dependent macrophages contribute to homeostatic control of the size of the liver. This study aimed to determine the systemic metabolic consequences of elevating circulating CSF1.

**Methods and Results:** Acute administration of a CSF1-Fc fusion protein led to monocytosis, increased resident tissue macrophages in the liver and all major organs, and liver growth. These effects were associated with increased hepatic glucose uptake and extensive mobilisation of body fat. The impacts of CSF1 on macrophage abundance, liver size and body composition were rapidly reversed to restore homeostasis. CSF1’s effects on metabolism were independent of several known endocrine regulators and did not impact the physiological fasting response. Analysis using implantable telemetry in metabolic cages revealed progressively reduced body temperature and physical activity with no change in diurnal food intake.

**Conclusion:** These results demonstrate the existence of a dynamic equilibrium between CSF1, the mononuclear phagocyte system, metabolic regulation and homeostatic control of liver:body weight ratio.

## 1.0 Introduction

The liver-body weight ratio in adults is maintained at a constant ratio to body weight [1]. Compensatory liver regeneration following partial hepatectomy involves an increased metabolic demand and global changes in metabolism, including reduced circulating glucose and lipid mobilisation from adipose, proportional to the degree of hepatic insufficiency [2, 3]. During the proliferative response in a growing liver a distinct subset of metabolically active cells appears to compensate for temporary deficits in liver function [4, 5].

The liver contains a large population of sinusoidal resident tissue macrophages, Kupffer cells, that are lost along with liver mass following partial hepatectomy. The major physiological macrophage growth factor, macrophage colony-stimulating factor (CSF1) is internalised and degraded by Kupffer cells, providing a homeostatic mechanism to control monocyte release from the bone marrow [6]. Consequently, circulating CSF1 concentrations rise in response to partial hepatectomy or liver injury, and subsequently fall in line with restoration of liver mass [7, 8]. Supporting a functional role for this response in regeneration, CSF1 treatment of mice accelerates liver regeneration following injury or resection [9, 10], and treatment of naïve mice, rats or pigs is sufficient to drive hepatocyte proliferation and an increase in the liver-body weight ratio[11-13].

Diverse tissue resident macrophage populations contribute to metabolic homeostasis, by regulating physiological mobilisation of fatty acids between adipose, liver and muscle and responses to insulin [14, 15]. In white adipose tissue (WAT), macrophages regulate both lipid storage and lipolysis at steady state and during obesity. Distinct roles have been attributed to resident and recruited macrophages[16, 17]. These observations led us to ask how CSF1R-dependent macrophages contribute directly or indirectly to meeting the metabolic demands of the growing liver and homeostatic maintenance of the liver-body weight ratio.

## 2.0 Methods

### 2.1 Animal Experiments

Animal studies utilised 1 mg/kg porcine (P)-CSF1-Fc [13] or 5 mg/kg Human CSF1-Mouse Fc conjugate (HM-CSF1-Fc, produced by Novartis, Switzerland), administered by sub-cutaneous injection, as indicated in results/figure legends. The two CSF1-Fc reagents were used interchangeably, based upon their availability. Previously reported minor differences in dose and precise time courses between the two CSF1-Fc reagents *in vivo* may be related to differential affinity for mouse CSF1R and/or impacts of the Fc region on circulating half-life [10]. The majority of studies were conducted in male mice on the C57BL/6J genetic background, unless otherwise indicated. Animals were housed in specific pathogen free facility on 12 hours dark cycle at 22-24°C with access to water and food ad libitum except during fasting. Fasting experiments were conducted from 8 am – 8 am (ZT1 in our animal facility). All animal experiments were approved by a University of Queensland Animal Ethics Committee.

Detailed methods are listed in Supplementary Information.

## 3.0 Results

### 3.1 CSF1-Fc treatment causes a transient increase in tissue macrophage populations and alters body composition

As reported previously [9, 13], daily CSF1-Fc treatment for 4 days led to an increase in liver and spleen size 24 h after the last treatment. Figure 1A-C show an extended time course of the changes in total body weight, liver and spleen weight demonstrating that this response is rapidly reversible; peaking on day 7 then declining over the following 7-10 days. To gain further insight into the increase in body weight and metabolic impacts in response to CSF1, we analysed body composition by NMR (Figure 1D-F). CSF1-Fc treatment led to a transient and rapidly reversible increase in lean mass and fluid mass and a 40% loss of fat, each of which was maximal on day 7. Consistent with previous findings in non-human primates and human patients infused with recombinant CSF1 [18, 19], short term CSF1-Fc treatment induced a profound monocytosis, which was also rapidly reversed; peaking on days 5-7 and returning to baseline by day 11 (Figure 1G).

**Figure 1.**
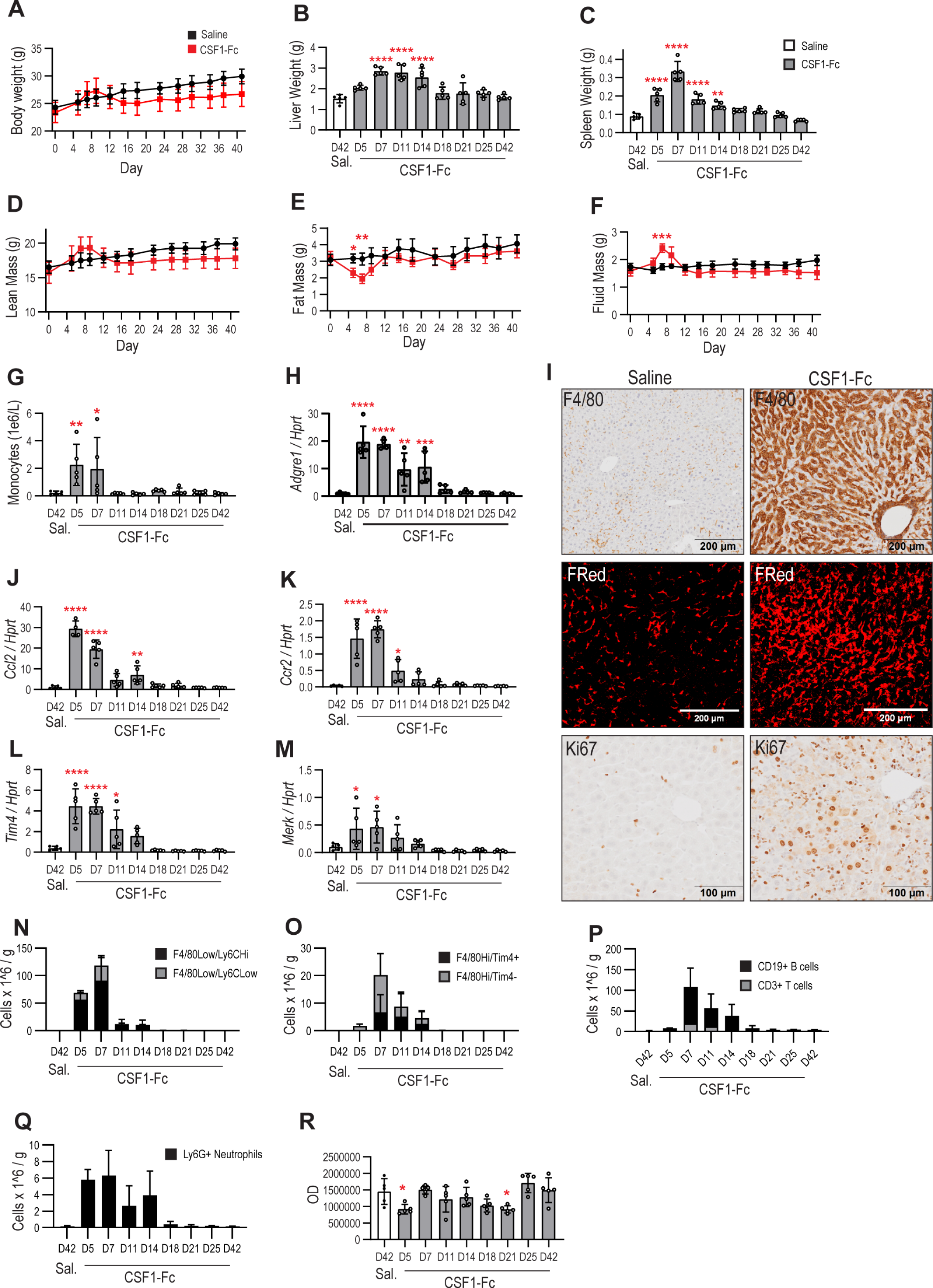
Acute CSF1-Fc treatment reversibly alters liver weight and body composition. Groups of 6-8 weeks old male mice (n=5/group) were administered 1 mg/kg P-CSF1-Fc for 4 days prior to sacrifice at the indicated timepoints up to 42 days. Control mice were treated with saline and sacrificed at 42 days. Wholemount images in panel I were generated from Csf1r-FusionRed mice treated with 5 mg/kg HM-CSF1-Fc for 4 days prior to sacrifice on day 5. Body weight (A), liver weight (B), spleen weight (C) lean mass (D), fat mass (E) and fluid mass (F) were measured at the indicated timepoints (Bruker Minispec). (G) Circulating blood monocytes. Liver *Adgre1* (H) mRNA expression, F4/80 staining, *Csf1r*-FusionRed expression and Ki67 staining (I) (representative images). Liver *Ccl2* (J), *Ccr2* (K), *Timd4* (L) and *Mertk* (M) expression. Liver monocyte (N) and macrophage (O) subsets, lymphocyte subsets (P) and neutrophils (Q) identified and quantified by flow cytometry. Caspase 3 enzyme activity in liver homogenates (R). Data are presented as mean +/- s.d. Two-way ANOVA with multiple comparisons (A, D-F); Ordinary one-way ANOVA with multiple comparison (B-C, G-H, J-M, R).

The transient increase in liver size was associated with increased Ki67 expression in both parenchymal and non-parenchymal cells, increased F4/80 staining (a marker for macrophages) and expression of the *Csf1r-*Fusion Red (FRed) reporter [20]. In parallel, increased mRNA expression of *Adgre1* (encoding F4/80), resident macrophage markers *Timd4* and *Mertk*, and both the monocyte chemoattractant *Ccl2* and its receptor *Ccr2* was observed (Figure 1H-M). Flow cytometric analysis of liver macrophage and monocyte sub-populations recovered following tissue disaggregation demonstrated a rapid transient increase in classical (F4/80^Low^/Ly6C^Hi^) and non-classical (F4/80^Low^/Ly6C^Low^) monocytes, as well as Kupffer cells (F4/80^Hi^/TIMD4^+^) and monocyte-derived macrophages (F4/80^Hi^/TIMD4^-^) (Figure 1L-M, gating strategy shown in Supplementary Figure 1). Hepatic granulocyte and lymphocyte content also increased transiently, likely reflecting CSF1-inducible chemokine expression[9] (Figure 1N-Q). There was no evidence of widespread apoptosis, measured by caspase 3 activity in liver homogenates, associated with the return to normal liver size post-treatment (Figure 1R).

We next asked whether the increase in tissue macrophages was restricted to the liver. CSF1-Fc treatment strikingly increased monocyte/macrophage abundance in both visceral and sub-cutaneous adipose, shown by F4/80 and *Csf1r-*FRed expression, with expression of *Adgre1, Ccr2* and *Ccl2* peaking at day 5, preceding maximal fat loss (day 7) (Figure 2A-E). Reduced fat mass and adipose tissue macrophage expansion was associated with a significant decrease in adipocyte size, and an increase in circulating triglyceride, peaking on day 5 (Figure 2 F,G). There was no significant change in the expression of *Pdgfc*, which was previously demonstrated to be a CSF1R-dependent gene expressed in adipose resident macrophages that promotes lipid storage[16] (Figure 2H). Like Kupffer cells in the liver, resident adipose tissue macrophages have a unique gene expression profile that includes constitutive expression of transcripts such as *Retnla, Mrc1, Clec10a* and *Arg1*, sometimes considered as markers of an anti-inflammatory phenotype[21], and meteorin-like (*Metrnl*), a hormone-like molecule that regulates thermogenesis[22]. CSF1-Fc treatment increased *Arg1, Clec10a, Mertk, Timd4* and *Metrnl* expression in adipose, indicating expansion and/or differentiation of resident macrophages, but had little impact on *Cd9* and *Lpl* (both expressed at high levels by adipocytes[23, 24]) Figure 2I-Q). Monocyte/macrophage populations were also increased in other key metabolic organs; including pancreas and skeletal muscle (Figure 2R,S). In line with liver and adipose, CSF1-Fc induced *Adgre1, Ccr2, Ccl2* and *Metrnl* in the tibialis anterior muscle (Figure 2T-W).

**Figure 2.**
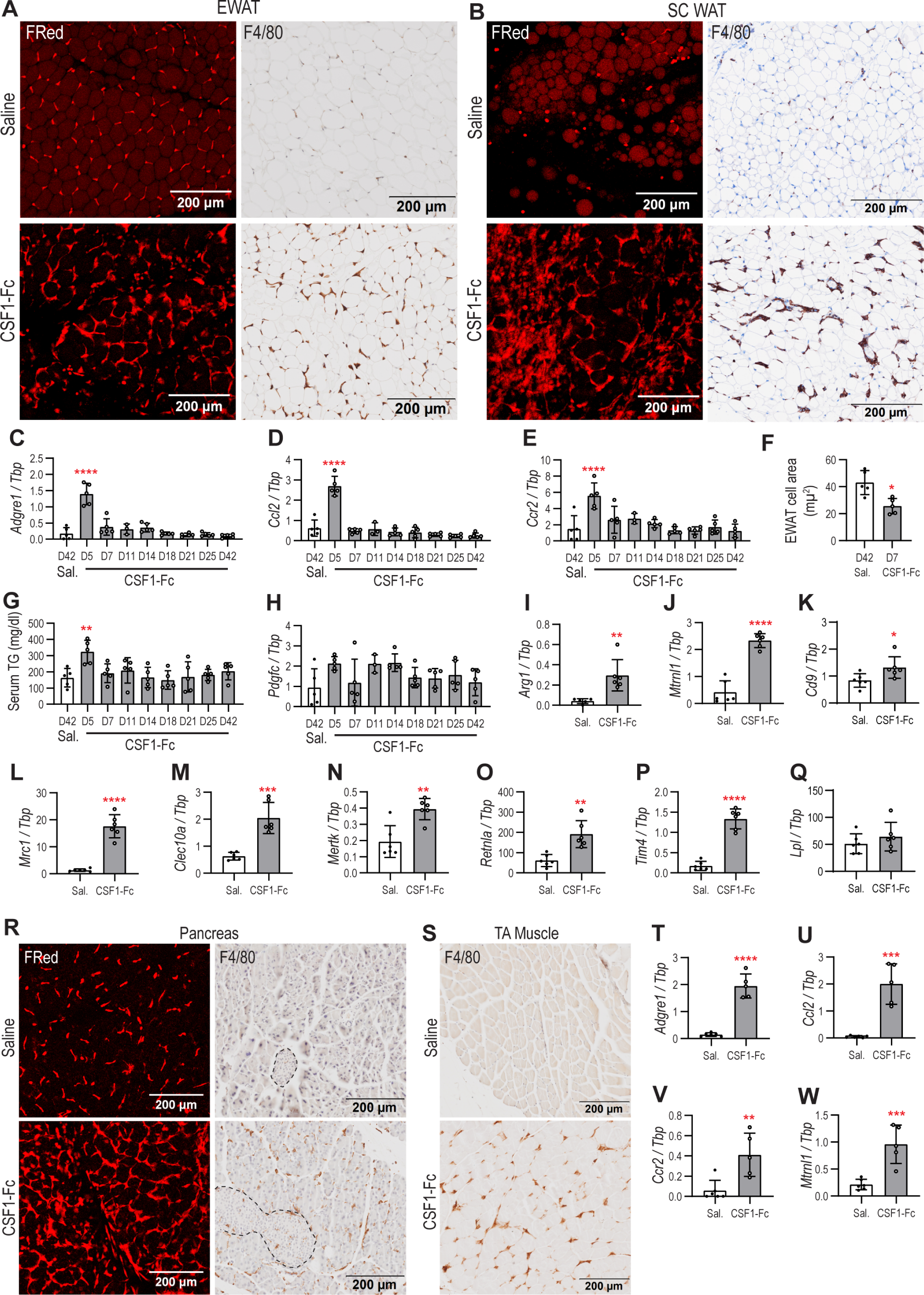
CSF1-Fc treatment induced macrophage expansion in adipose tissue and reduced adipocyte size. Groups of male mice (n=5/group) were administered 1 mg/kg P-CSF1-Fc for 4 days prior to sacrifice at the indicated timepoints up to 42 days. Control mice were treated with saline and sacrificed at 42 days. Wholemount images in panels A, B and R, and qPCR in panels I-Q and T-W were generated from Csf1r-FusionRed mice treated with 5 mg/kg HM-CSF1-Fc for 4 days prior to sacrifice on day 5 (n=6/group). Adipose macrophages detected by direct imaging of Csf1r-FusionRed or F4/80 IHC in epidymal (EWAT) (A) and sub-cutaneous (SC WAT) (B) white adipose tissue (representative images). Adipose *Adgre1* (C) *Ccl2* (D) and *Ccr2* (E) mRNA expression. (F) EWAT adipocyte area. Serum triglyceride concentration (G). Adipose *Pdgfc* (H), Arg1 (I), Mtrnl (J), Cd9 (K), Mrc1 (L), Clec10a (M), Mertk (N), Retnl (O), Timd4 (P) and Lpl (Q) mRNA expression. Pancreas (R) and tibialis anterior (TA) muscle macrophages (S) detected by direct imaging Csf1r-FusionRed or F4/80 IHC (representative images). TA muscle Adgre1 (T), Ccl2 (U), Ccr2 (V) and Mtrnl (W) mRNA expression. Data are presented as mean +/- s.d. Ordinary one-way ANOVA with multiple comparison (C-E, G-H); Two-tailed unpaired t test (F, I-Q, T-W).

### 3.2 CSF1-Fc treatment alters glucose metabolism

Partial hepatectomy in mice leads to a fall in glucose and mobilisation of lipid from adipose to the liver [25]. CSF1-Fc treatment caused a transient loss of circulating insulin and insulin-like growth factor 1 (IGF1) associated with reduced hepatic *Igf1* mRNA expression, which is the main source of circulating IGF1 (Figure 3A-C). A significant decline in random blood glucose was associated with reduced circulating insulin and IGF1 (Figure 3D-F) and an increase in circulating glucagon (Figure 3G). To determine the mechanism behind the decline in blood glucose, mice were injected with ^18^F-Fluorodeoxyglucose (^18^F-FDG) after 1 or 4 daily injections of CSF1-Fc and the distribution of label was assessed by Positron Emission Tomography and Computed Tomography (PET-CT). After 4 days of treatment with CSF1-Fc there was a massive and selective increase in hepatic uptake of ^18^F-FDG (Figure 3H-K). There was also evidence of uptake in the spleen, which was also rapidly growing in response to CSF1-Fc (Figure 3I). In hepatocytes FDG uptake is reversible and it does not normally accumulate because of the activity of glucose-6-phosphatase. Indeed, FDG imaging has been used in mice to monitor increased uptake of glucose by inflammatory lymphocytes in hepatitis models [26]. Thus, hepatic FDG accumulation may be due in part to glucose uptake by macrophages themselves. Despite the hepatic glucose uptake, hepatocyte glycogen content was significantly reduced in CSF1-Fc-treated mice, with the nadir coinciding with peak liver growth (Figure 3L,M).

**Figure 3.**
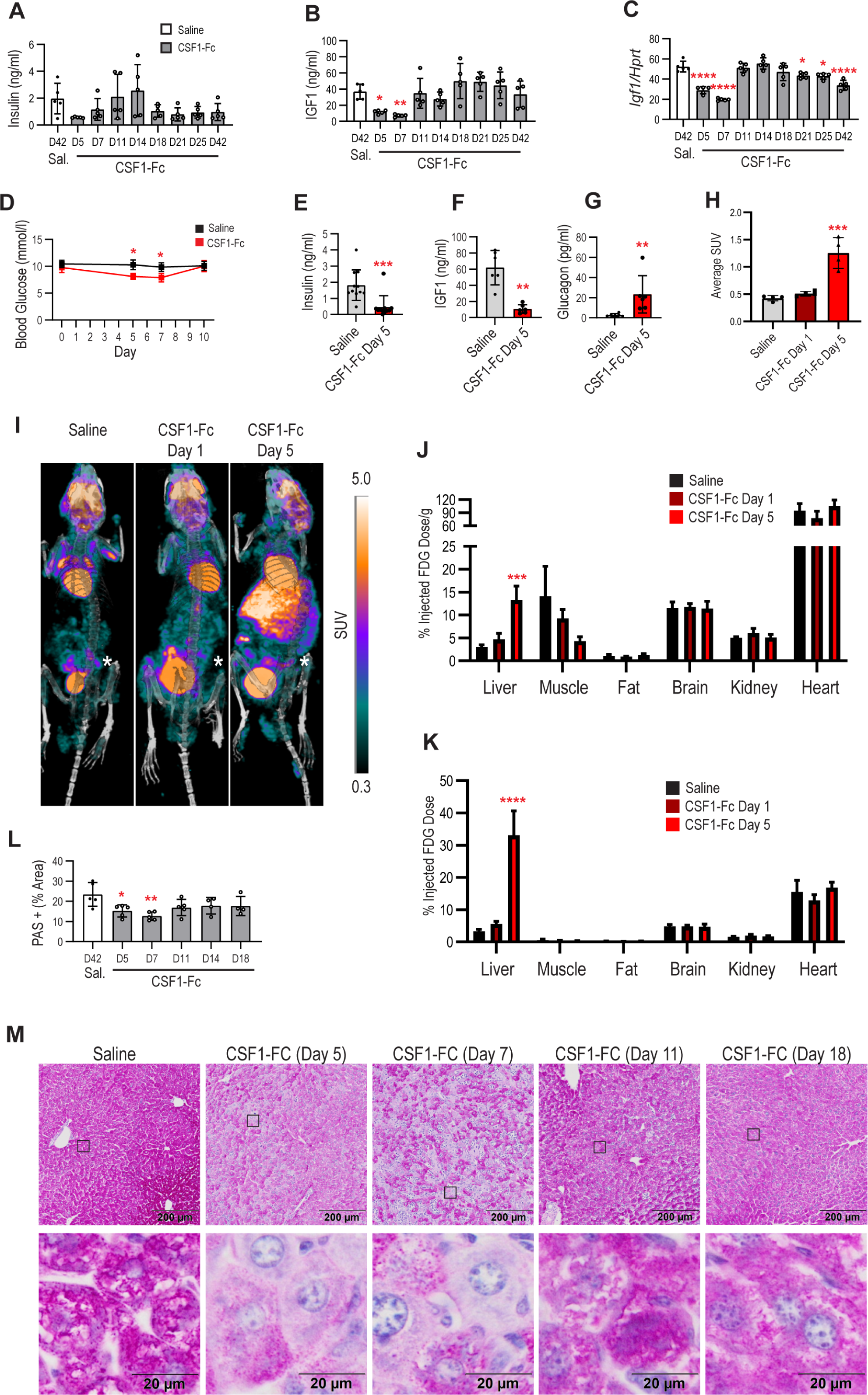
CSF1-Fc treatment reduced circulating IGF1 and induced hepatic glucose uptake. Groups of male mice (n=5/group) were administered 1 mg/kg P-CSF1-Fc for 4 days prior to sacrifice at the indicated timepoints up to 42 days. Control mice were treated with saline and sacrificed at 42 days. Serum insulin (A) and IGF1 (B) were measured by ELISA, and liver *Igf1* expression (C) was measured by qPCR. Groups of 6 male mice were treated for 4 days with saline control or 5 mg/kg HM-CSF1-Fc and sacrificed on day 5 or day 10. Blood glucose (D) was measured using a glucometer and insulin (E), IGF1 (F) and glucagon (G) were measured by ELISA. For the analysis of glucose uptake, groups of 4 male mice were treated with 5 mg/kg HM-CSF1-Fc for 1 day or 4 days prior to PET-CT imaging of [^18^F]Fluorodeoxyglucose (FDG) uptake (representative images in I, with the level of uptake of labelled FDG represented as Standardised Uptake Value (SUV). Asterisk indicates the location of the spleen). (H) Average SUV across the whole image for the 4 animals at each time point. The % injected FDG dose (J) and % dose/g tissue (K) in dissected tissues were determined using a gamma counter. Liver glycogen quantified by Periodic Acid Schiff (PAS) staining area (L (n=4/group, subset of animals and timepoints from (A)), representative images in (M)). Data are presented as mean +/- s.d. Ordinary one-way ANOVA with multiple comparison (A-C, F, G, H-K, L); Two-way ANOVA with multiple comparisons (D); Two-tailed unpaired t test (E-G).

C57BL/6J mice have a null mutation in the nicotinamide nucleotide transhydrogenase (*Nnt*) gene which impacts many aspects of metabolism including adiposity and glucocorticoid production (reviewed in [27]). We therefore tested whether the CSF1-Fc response was mouse strain-specific. The effects of CSF1-Fc treatment on body composition, organ weights, monocyte-macrophage mobilisation and blood glucose were also observed in Balb/c mice (*Nnt* wildtype, among many other genetic variations) (Supplementary Figure 2).

### 3.3 Neither CSF1-stimulated macrophage expansion nor anti-CSF1R-mediated macrophage depletion affects the response to fasting

Regenerating liver maintains physiological responses to metabolic demand [4, 5]. The metabolic response to short-term fasting in mice has been widely studied with well-documented impacts on hormone balance, body weight, metabolism and hepatic enzymes [28]. Fasting leads to adipose tissue lipolysis, largely mediated by catecholamine signalling, and release of free fatty acids (FFA) and glycerol which are taken up by the liver [29, 30]. We next asked whether CSF1R-dependent macrophages regulate the response to fasting. As expected, 24 h fasting led to a significant loss of lean body mass, fat and fluid. This response was not altered by tissue macrophage expansion induced by CSF1-Fc pre-treatment (Figure 4A-D). We observed a reduction in blood monocytes in response to 24 h fasting, which has previously been reported to be glucocorticoid dependent [31], which reached significance in CSF1-Fc pre-treated mice (Figure 4E). The effects of CSF1-Fc treatment and fasting on body composition were essentially additive, so that the combined treatment led to the loss of 70-80% of total body fat. Fasting led to a small reduction in the weight of the liver but did not completely reverse the CSF1-Fc-stimulated increase in liver size (Figure 4F). CSF1-Fc produced a marginal reduction in glycogen in the fed state but did not prevent further loss of glycogen upon fasting (Figure 4G).

**Figure 4.**
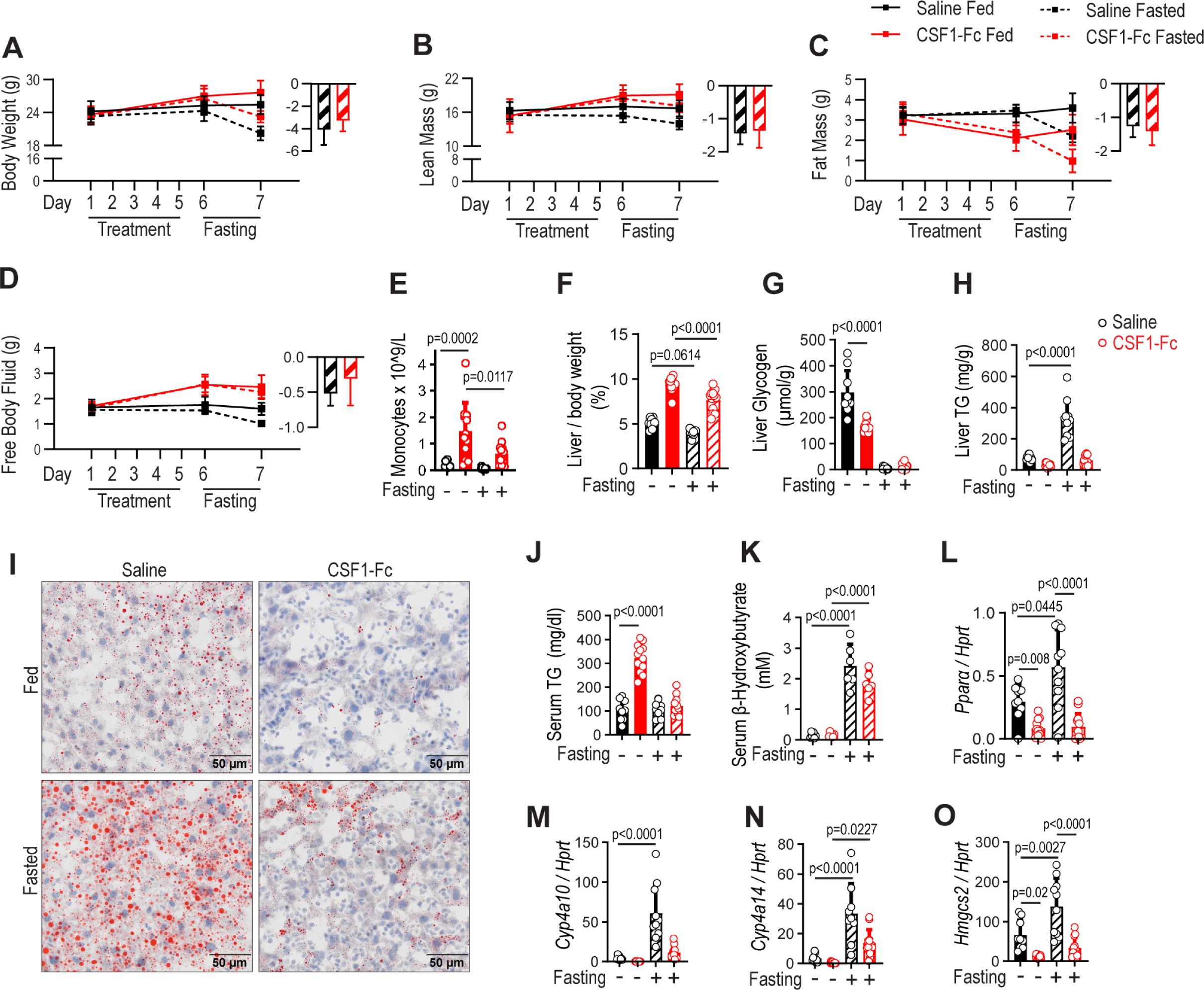
CSF1-Fc treatment does not impact the metabolic response to fasting. Groups of 6 male mice were treated with 1 mg/kg P-CSF1-Fc or saline control for 4 days, fasted for 24 h commencing on day 6 and sacrificed on day 7 (data from replicate experiments were combined for total n=12). Body weight (A), lean mass (B), fat mass (C) and free body fluid (D) were measured in each mouse pre-treatment and pre- and post-fasting. Right hand sub-panels in A-D show the average change from the pre-fasting level (day 6) in each parameter following 24 hrs fasting. These changes were significant for body weight and fat mass (p<0.0001 for both saline and CSF1-Fc treated) and for free body fluid (p=0.0002 for saline, p=0.0237 for CSF1-Fc treated). There were no significant changes in any of these parameters in the unfasted cohort. Blood monocytes (E), liver weight (F), glycogen (G) and triglyceride (TG) content (H) were measured post-sacrifice. Representative Oil Red O lipid staining (I). Serum TG (J) and B-hydroxybutyrate (K). Liver *Ppara* (L) *Cyp4a10* (M) *Cyp4a14* (N) and *Hmgcs2* (O) mRNA expression. Data are presented as mean +/- s.d. Two-way ANOVA with Sidak’s test for multiple comparisons.

CSF1-Fc treatment reduced hepatic triglyceride in the fed state and prevented lipid accumulation in response to fasting (Figure 4H,I). By contrast, CSF1-Fc treatment induced a 3-fold increase in serum triglycerides, which was almost completely reversed in response to fasting (Figure 4J). Despite the lack of hepatic lipid accumulation, CSF1-Fc pre-treatment did not affect circulating β-hydroxybutyrate levels indicating that normal fasting-induced ketogenesis was neither impaired nor augmented (Figure 4K). The liver response to fasting involves the activation of PPARα. Fasting caused marginal up-regulation of *Ppara* mRNA expression, but the PPARα target genes *Cyp4a10* and *Cyp4a14* [32] were strongly induced. Prior CSF1-Fc treatment damped the expression of these PPAR target genes in both fed and fasting mice likely reflecting the reduction in triglyceride accumulation in both states (Figure 4L-N). CSF1-Fc also downregulated expression of *Hmgcs2*, encoding HMGCS, the rate-limiting enzyme in β-hydroxybutyrate synthesis (Figure 4O).

To further investigate the physiological impact of the expansion of tissue macrophage populations metabolomic profiling was performed on serum and liver from fed and fasted mice (Table S1). We utilised the network analysis tool Graphia to analyse clusters of correlated metabolites. Fed and fasted samples were inter-correlated in both serum and liver, irrespective of prior CSF1-Fc treatment (Figure 5A,E). Serum metabolites clustered into groups of metabolites that were depleted (Cluster 1, including many sugars) or enriched (Cluster 2, including hydroxybutyric acid) in fasted mice. There was no clear cluster of CSF1-Fc-regulated metabolites (Figure 5B, Table S1). The metabolomic analysis detected the decrease in blood glucose in response to both CSF1-Fc treatment and fasting (Figure 5C). The most highly CSF1-Fc-regulated metabolite in serum was 5-methoxytrytamine (Figure 5D), a tryptamine derivative related to the neurotransmitters serotonin and melatonin. Despite the reduction of total body fat mass in response to both fasting and CSF1-Fc treatment, and the detected increase in triglycerides, neither fasting nor CSF1-Fc had any significant effect on circulating free fatty acids, glycerol or cholesterol (Table S1).

**Figure 5.**
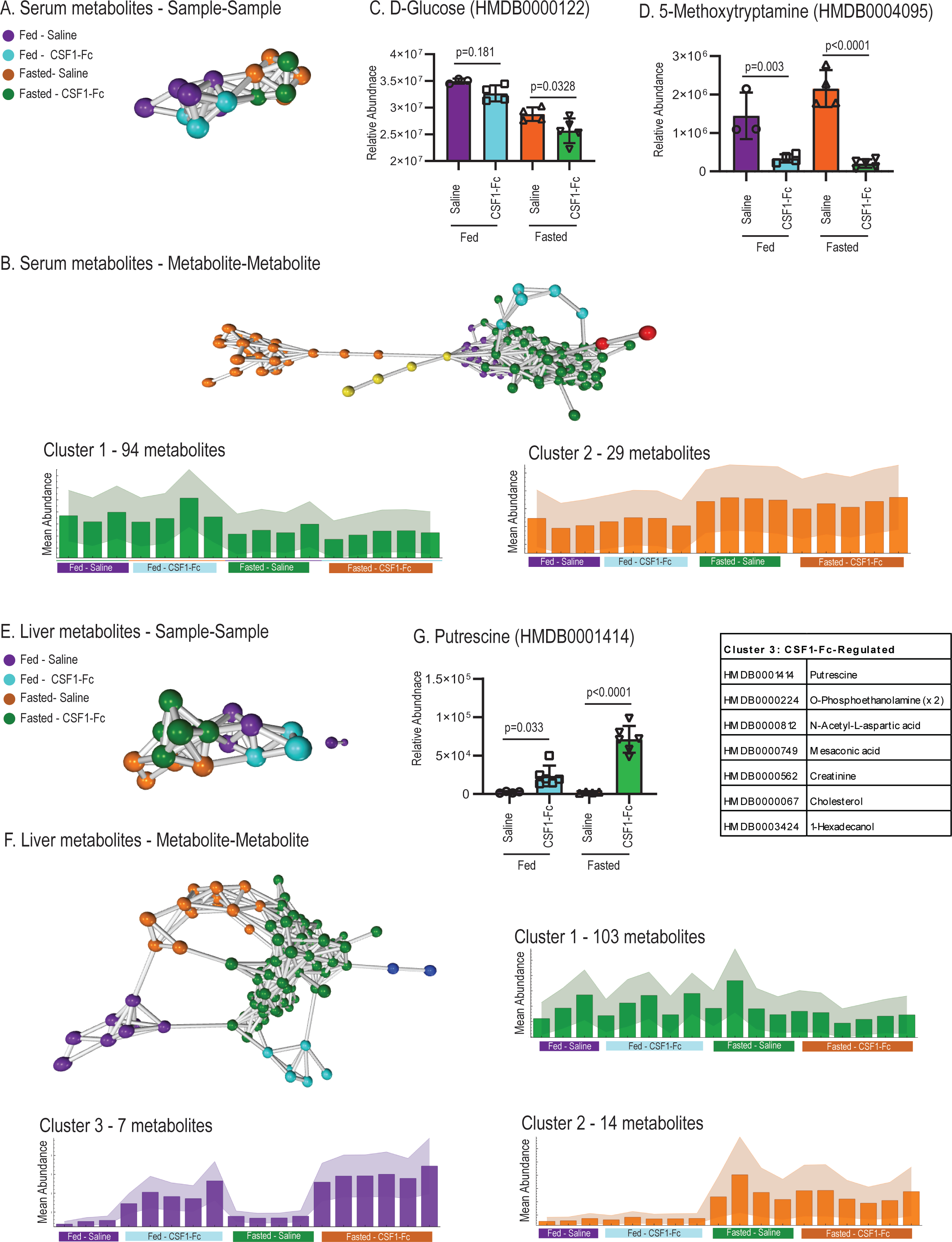
The impact of fasting and CSF1-Fc treatment on serum and liver metabolites. Groups of 6 male mice were treated with 1 mg/kg P-CSF1-Fc or saline control for 4 days, fasted for 24 h commencing on day 6 and sacrificed on day 7. Gas chromatography-Mass spectrometry-based metabolomics was performed on serum and liver homogenates. Relative quantification data (MS peak area) was analysed in Graphia. Serum sample-sample (A) and metabolite-metabolite clustering (B). Relative abundance of serum glucose (C) and 5-methoxytryptamine (D). Liver sample-sample (D) and metabolite-metabolite clustering (E). Relative abundance of liver putrescine (G). CSF1-Fc-regulated liver metabolites comprising cluster 3 (H). Clusters with fewer than 3 metabolites not shown. Histograms show mean of all metabolites in indicated clusters, with area shading indicating minimum/maximum standard error. C,D,G: data are presented as mean +/- s.d. One way ANOVA, Sidak’s multiple comparisons test.

The dominant impact of fasting on liver metabolites was a reduction in phosphorylated sugars; glucose-6-phosphate, mannose-6-phosphate, fructose-6-phosphate and fructose-1-phosphate, indicative of the switch from glycolysis to gluconeogenesis, which was unaffected by CSF1-Fc treatment (Figure 5F, Cluster 2, Table S1). There was a small cluster of CSF1-Fc-regulated metabolites that included putrescine and phosphoethanoloamine, which likely reflect liver growth (Figure 5F-H, cluster 3, Table S1). Polyamines have well-documented functions in cell proliferation and putrescine levels were reported to increase up to 50-fold in regenerating rat liver [33]. Taken together these findings indicate that lipolysis and release of free fatty acids in response to fasting is balanced by their metabolism in tissues and the production of ketone bodies.

Loft *et al.* [34] reported that induction of ketogenesis in response to fasting was dependent in part upon glucocorticoid signalling in resident hepatic macrophages. To test this conclusion, we performed the reciprocal experiment, to determine whether resident macrophages were required in the lipolytic response to fasting. Groups of mice were treated twice weekly with the anti-CSF1R antibody AFS98 for 2 weeks, followed by a 24 h fast initiated 3 days after the final dose. The efficacy of CSF1R blockade was demonstrated by the large increase in circulating CSF1 in AFS98-treated mice (Supplementary Figure 3A). Anti-CSF1R treatment completely ablated the abundant *Csf1r*-FRed-positive resident macrophages in most tissues examined, notably adipose, pancreas, kidney and smooth muscle (Supplementary Figure 3B). Partial macrophage depletion was achieved in the liver; quantified by assessing the loss of *Adgre1* mRNA (Supplementary Figure 3B,C). Hepatic *Ccl2* mRNA expression was not affected by AFS98 treatment but was reduced to almost undetectable levels by fasting (Supplementary Figure 3D). Anti-CSF1R treatment did not alter the trajectory of physiological body weight gain but did cause a small reduction in liver:body weight ratio (Supplementary Figure 3E,F) as observed previously following more prolonged anti-CSF1R treatment [35]. There was no effect of global macrophage depletion on body composition, blood glucose or liver weight reduction in response to fasting, although AFS98-treated mice had a modest reduction in adipocyte size, which was significant in the fed state. Global macrophage depletion did not impact the fasting-induced accumulation of liver lipid, the induction of PPARα target gene expression, or circulating β-hydroxybutyrate (Supplementary Figure 3G-O).

### 3.4 The response to CSF1-Fc is not affected by blockade of ***β***-adrenergic signalling or IGF1 supplementation

The catecholamine/β-adrenergic signalling axis is an essential hormonal system regulating body composition and metabolism. To support the conclusion that the CSF1-Fc response is independent of known physiological mechanisms of metabolic regulation, mice were pre-treated with the non-selective β-adrenergic receptor inhibitor propranolol (10 mg/kg [36]) on day −1, followed by daily co-administration of propranolol and CSF1-Fc or saline control for 4 days and sacrifice on day 5. Inhibition of β-adrenergic signalling with propranolol did not prevent the CSF1-Fc-induced alterations in body composition, liver and spleen growth or monocytosis (Figure 6A-G).

**Figure 6.**
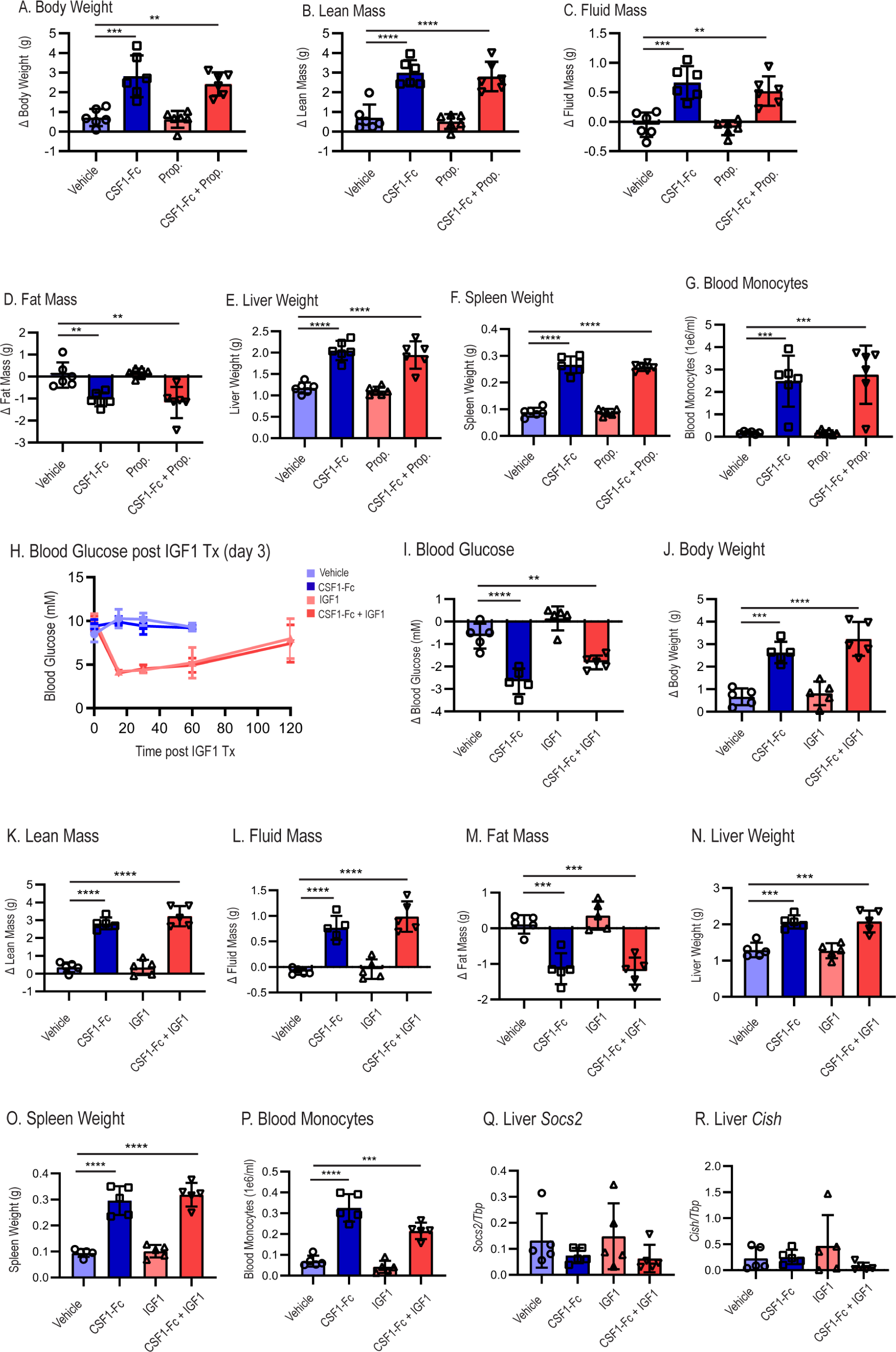
Blockade of β-adrenergic signalling or IGF1 treatment do not impact the response to CSF1-Fc. (A-G) Groups of male mice were treated with propranolol or saline control (10 mg/kg) one day prior to commencing CSF1-Fc treatment, then co-treated with propranolol and/or CSF1-Fc (5 mg/kg HM-CSF1-Fc) or saline control daily by sub-cutaneous injection for 4 days and sacrificed on day 5. Body weight (A), lean mass (B), fluid mass (C), fat mass (D), liver weight (E), spleen weight (F) and blood monocyte count (G) at sacrifice. (H-R). Groups of male mice were treated once daily with CSF1-Fc (5 mg/kg HM-CSF1-Fc) or saline control by subcutaneous injection and/or IGF1 (5 mg/kg, split into 2 doses) or saline control by intraperitoneal injection for 4 days and sacrificed on day 5. Blood glucose levels following IGF1 treatment (day 3 of CSF1-Fc treatment) (H). Blood glucose (I), body weight (J), lean mass (K), fluid mass (L), fat mass (M), liver weight (N), spleen weight (O), blood monocyte count (P), liver *Socs2* (Q) and *Cish* (R) mRNA expression at sacrifice. Data are presented as mean +/- s.d. Ordinary one-way ANOVA with multiple comparisons.

CSF1-Fc treatment led to a substantial reduction in both insulin and IGF1 in the circulation (Figure 2E,F). Insulin and IGF1 are closely-related and IGF1 can bind to both the insulin receptor (INSR) and IGF1R [37]. IGF1 produced by adipocytes and macrophages has been reported to contribute to adipose homeostasis and both insulin and IGF1 can suppress lipolysis [38]. To address the potential importance of reduced insulin/IGF1, CSF1-Fc was co-administered with recombinant human IGF1 (5 mg/kg/day). As anticipated, IGF1 administration caused sustained hypoglycaemia (Figure 6H/I), confirming the efficacy of the treatment, but there was no effect on the gross systemic response to CSF1-Fc, including liver weight, fat mass, circulating monocytes and glucose (Figure 6J-P). This outcome indicates that the fall in blood glucose *per se* does not mediate the effects on fat mobilisation. Reduced circulating IGF1 could indirectly lead to increased pituitary expression of growth hormone (GH). However, GH-signalling in the liver, assessed by analysis of GH target gene expression (*Socs2, Cish*), was unaffected (Figure 6Q,R).

### 3.5 IL6 is not required for the response to CSF1-Fc

Interleukin 6 (IL6) is a known hormone-like regulator of metabolism, tissue regeneration and adipose homeostasis [39] and hepatic *Il6* mRNA was increased in CSF1-Fc-treated mice [9, 13]. A time analysis of this response showed a 50-fold increase in *Il6* mRNA in the liver which remained elevated until day 14 (Supplementary Figure 4A). This was not a classical acute phase response; serum amyloid precursor (*Apcs*) mRNA was increased only marginally and serum amyloid A1 (*Saa1*) was almost completely repressed (Supplementary Figure 4B,C). In CSF1-Fc treated mice, serum IL6 was increased 6-fold on day 7, coinciding with the peak CSF1-Fc response, and was still significantly elevated at day 11 (Supplementary Figure 4D). To test the role of IL6 as a downstream mediator of the CSF1-Fc response mice were treated with anti-IL6 antibody prior to (day −1) and during (day 2) a 4-day CSF1-Fc treatment regime. The upregulation of circulating IL6 and significant neutralisation by anti-IL6 was confirmed (Supplementary Figure 4E). IL6 blockade did not alter the response to CSF1-Fc in terms of monocytosis, increased liver or spleen weight, change in body composition, blood glucose or the loss of circulating IGF1 (Supplementary Figure 4F-N).

### 3.6 CSF1-Fc-induced fat mobilisation occurs in the absence of TYROBP signalling

The macrophage-expressed lipid sensor TREM2 regulates lipid metabolism in liver and adipose tissue, especially in response to excess lipid [40, 41]. TREM2 and its adaptor protein TYROBP synergise with CSF1R to activate downstream pathways that regulate macrophage proliferation and function [42]. Liver and adipose expression levels of *Trem2* and *Tyrobp* were markedly elevated in response to CSF1-Fc treatment (Supplementary Figure 5A-D). To test the hypothesis that TREM2/TYROBP signalling may facilitate the lipid mobilisation observed in CSF1-Fc treated mice, a novel TYROBP knockout (KO) mouse line was created using CRISPR/Cas9 gene editing (Materials and Methods). Bone marrow cells from the TYROBP KO line exhibited an impaired response to CSF1, as reported for the original knockout [42], but differentiated normally in response to CSF2 (Supplementary Figure 5E). To investigate the role of TYROBP in the CSF1-Fc response *in vivo*, wild type and TYROBP KO mice were treated with CSF1-Fc or saline control for 4 days and sacrificed on day 5. No differences in CSF1-Fc induced monocytosis, effects on body composition, blood glucose, liver or spleen growth were observed between WT and KO mice (Supplementary Figure 5F-L). CSF1-Fc treatment induced liver *Trem2* and *Tyrobp* mRNA expression equally in both WT and KO mice (Supplementary Figure 5M,N).

### 3.7 The rapid growth response to CSF1-Fc treatment is associated with increased fat utilisation and transient reduction in body temperature and activity

To further investigate the mechanism of fat mobilisation in response to CSF1-Fc treatment, mice were housed individually in metabolic cages with implanted telemetry to continuously monitor weight, food and water intake, body temperature, physical activity and metabolism. CSF1-Fc treatment did not affect food or water intake, during treatment or recovery, in absolute terms or per g body weight (Figure 7A-B). At the time of peak fat reduction (day 5), which preceded the peak increase in body weight, there was a small and transient reduction in respiratory exchange ratio (RER), suggestive of increased lipid utilisation (Figure 7C-E). Weight gain in CSF1-Fc-treated mice (day 3 to day 6) was associated with significantly reduced activity (Figure 8A), and reduced core temperature and resting metabolic rate on day 5 (Figure 8B-D). These data eliminate the possibility that CSF1-Fc acts via inappetence or increased physical activity. Given the surprising reduction in body temperature in response to CSF1-Fc, we examined brown adipose tissue (BAT). CSF1-Fc treatment increased BAT monocyte/macrophage content on day 5 post treatment commencement, as shown by increases in *Adgre1, Ccr2* and *Ccl2* mRNA expression and F4/80 staining (Figure 8E-H). Corroborating the reduction in core temperature, uncoupling protein 1 (*Ucp1*) mRNA expression in brown adipose tissue was significantly reduced in CSF1-Fc-treated mice (Figure 8I), indicating that mobilised lipid is not re-directed to thermogenesis.

**Figure 7.**
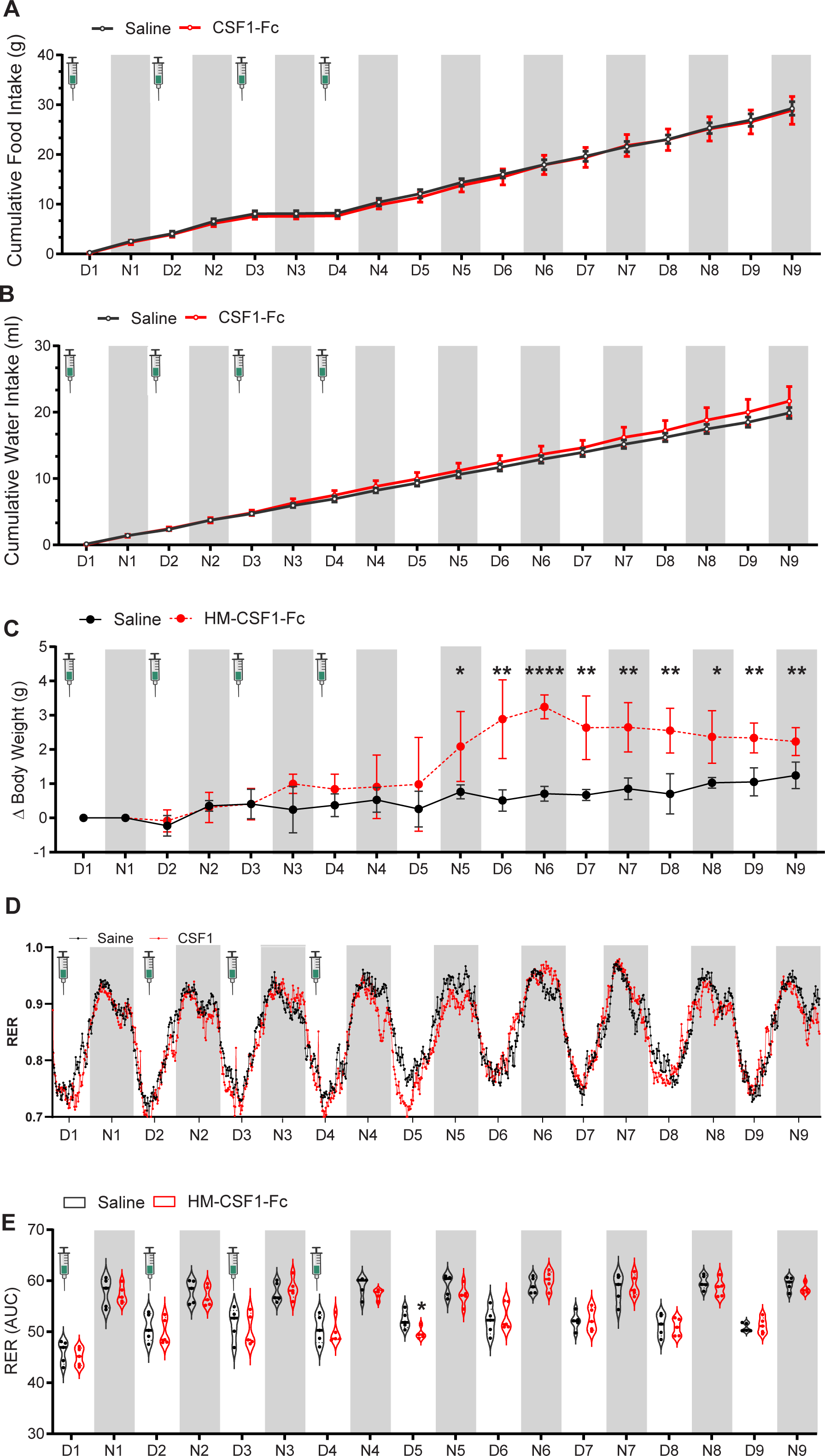
CSF1-Fc treatment increases body weight without altering food intake. Male mice with implantable telemetry devices were housed in metabolic cages (Phenomaster) for 7 days prior to treatment to acclimatise and maintained on a 12:12 light:dark cycle as indicated (D = day, N =night). Groups of 6 mice were then treated with CSF1-Fc (5 mg/kg HM-CSF1-Fc) or saline control by sub-cutaneous injection daily on 4 successive days at the times indicated and monitored in the metabolic cages until sacrifice on day 10. Cumulative food (A) and water (B) intake and change in body weight (C) during the experiment. Average respiratory exchange rate (RER) (D) and area under the RER curve (E). Data in A-C are presented as mean +/- s.d, D shows mean, E shows median (horizontal line) and distribution. Multiple unpaired student’s t-test with Welch correction comparing saline vs CSF1-Fc treatment (C,E).

**Figure 8.**
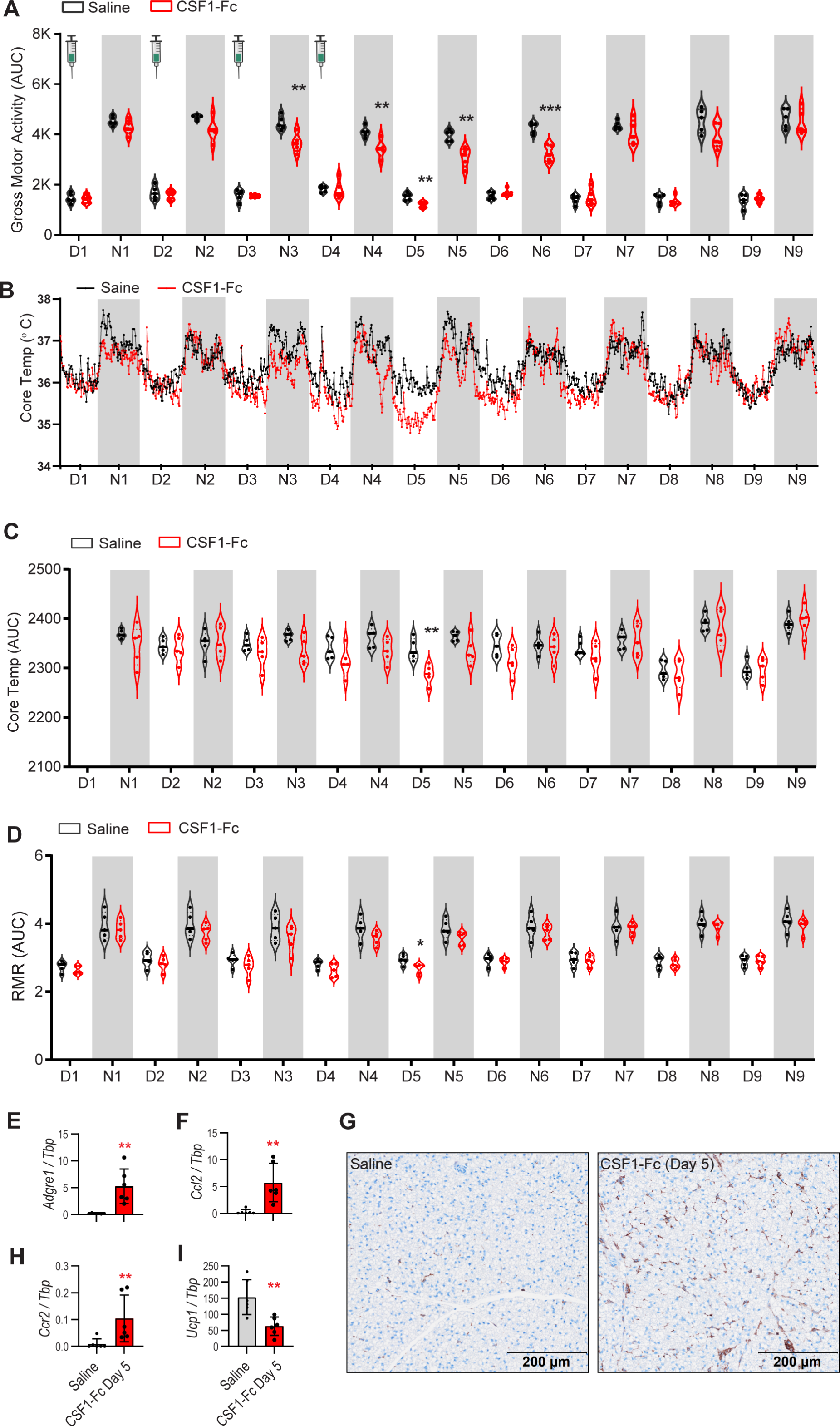
CSF1-Fc treatment decreases activity and core temperature. Male mice with implanted telemetry devices were housed in metabolic cages (Phenomaster) for 7 days prior to treatment to acclimatise and maintained on a 12:12 light:dark cycle as indicated (D = day, N =night). Groups of 6 mice were then treated with CSF1-Fc (5 mg/kg HM-CSF1-Fc) or saline control by sub-cutaneous injection daily on 4 successive days at the times indicated and monitored in the metabolic cages until sacrifice on day 10. Gross motor activity (A, Area Under the Curve (AUC)), core temperature (B and C) and resting metabolic rate (RMR, D) during the treatment period are shown. A separate cohort of mice was treated with CSF1-Fc (5 mg/kg HM-CSF1-Fc) or saline control by sub-cutaneous injection daily for 4 days and sacrificed on day 5 (n=6/group). Expression of *Adgre1* (E), *Ccl2* (F) and *Ccr2* mRNA in brown adipose tissue was quantified by qPCR. (G) Representative F4/80 staining in brown adipose tissue. (I) *Ucp1* mRNA expression in brown adipose tissue was quantified by qPCR. A, C and D show median (horizontal line) and distribution, B shows mean, E-F and H-I show mean +/- s.d. Multiple unpaired student’s t-test with Welch correction comparing saline vs CSF1-Fc treatment (A, C-D); Two-tailed unpaired t test (E-H).

## 4.0 Discussion

This study addresses the roles of CSF1 and CSF1R-dependent macrophages in meeting the metabolic demands of liver growth. The absolute increment in liver weight in response to CSF1-Fc is similar to that associated with recovery following 70% hepatectomy. We showed the systemic increase in tissue macrophages following CSF1-Fc administration is associated with metabolic alterations: reduced circulating glucose, hepatic glucose uptake and fat mobilisation. CSF1-Fc-induced liver growth shares commonalities with direct hyperplasia, in particular the lack of IL6 dependence, but the metabolic changes, including lipid mobilisation, resemble those observed in compensatory regeneration. In both surgical and toxic injury-induced regeneration, however, proliferative hepatocytes originate in the mid-lobular region, enabling portal and central hepatocytes to maintain their specialised functions, [4, 43], whereas hepatocellular proliferation in response to CSF1-Fc treatment was not zone restricted.

Heterozygous mice with a dominant kinase-dead mutation in the *Csf1r* gene do not respond to CSF1-Fc treatment, demonstrating that the response depends on CSF1R [44], which in turn is expressed only in macrophage lineage cells [20]. The postnatal growth and development of the liver is compromised in *Csf1r* knockout rats [45] and in CSF1-deficient mice (*Csf1*^op/op^) liver regeneration is delayed [46]. Our results further underline the importance of CSF1 in the so-called hepatostat; demonstrating that increased circulating CSF1 is sufficient to drive changes in metabolism to support growth of the liver and critically that the response is intrinsically reversible. Since circulating CSF1 is internalised and degraded by CSF1R-mediated endocytosis [6] increased macrophage numbers cannot be maintained without sustained CSF1 elevation. In essence, the maintenance of a constant liver:body weight ratio (the hepatostat) is linked to the CSF1-controlled dynamic equilibrium of the mononuclear phagocyte system.

Just as metabolic regulation is preserved during liver regeneration, the metabolic impacts of CSF1-Fc treatment were independent of known hormonal pathways regulating metabolism. The fall in blood glucose in response to CSF1-Fc treatment provided one plausible explanation for the fall in IGF1 and insulin in the circulation. Like insulin, IGF1 inhibits lipolysis in adipose tissue and Jordan *et al.* [47] reported the almost complete loss of *Igf1* mRNA in fasted liver. However, we found that co-administration of IGF1 with CSF1-Fc, to restore circulating IGF1 levels, had the expected impact on blood glucose but there was no effect on CSF1-Fc-induced fat loss or other systemic responses. The independence of the CSF1-Fc response from classic pathways of hormonal control was further supported by the lack of effect of co-administration of the β-blocker propranolol. Moreover, β-adrenergic activation increases thermogenesis [48], the opposite of the reduced body temperature observed here.

Maintaining body weight requires balancing energy from food intake with energy expenditure, while increasing weight requires that energy in food intake exceeds expenditure. No significant changes in food or water intake were observed in CSF1-Fc-treated mice. Reduced dark-phase activity was observed in treated mice 2 days after commencement of treatment and persisted for 3 days following treatment cessation, coinciding with a reduction in core body temperature and respiratory exchange ratio (RER), indicative of a shift in substrate utilisation from carbohydrates to fat [49]. These metabolic changes preceded body mass increase, which is largely attributed to liver growth, suggesting conservation of energy and mobilisation of fat to fuel growth. Reduced physical activity and energy generation from fat catabolism are also hallmarks of the energy conservation response to reduced food supply and cold, known as torpor when core temperature falls below 31°C. However, unlike the CSF1 response, torpor is associated with reduced food intake and increased ketogenesis [50].

Adipose resident macrophages and recruited monocytes have been attributed distinct functions in lipid storage and inflammation. Macrophage-produced PDGFcc was strongly implicated in promoting lipid storage and shown to be expressed in a CSF1R-dependent manner[16]. In contrast, Chen *et al* identified a role for CD169+ adipose resident macrophages in adipose lipid storage. Depletion of CD169+, but not CSF1R-dependent, macrophages on a normal chow diet reduced fat pad weight and induced adipocyte hypertrophy; and exacerbated pathological lipid accumulation in high fat diet-fed mice [17]. In the present study, acute expansion of adipose monocytes and macrophages where lipids are not in excess, led to fat mobilisation, without ectopic lipid accumulation, and decreased activity and core temperature. Our data suggest that PDGFcc may not be sufficient to promote lipid storage as expression remained high in adipose from CSF1-Fc treated mice. Consistent with the lack of visceral adipose in *Csf1r* knockout rats [51], macrophage or PDGFcc deficiency prevented postnatal adipose development in mice [16], which may suggest different roles for macrophages in neonates and adults.

CSF1-Fc treatment massively increased *Trem2* and *Tyrobp* mRNA in the liver and adipose tissue. Although TYROBP acts as an adaptor for several receptors, TREM2 is the most widely studied, as mutations in both TREM2 and TYROBP are strongly linked to Alzheimer’s disease risk [52]. TYROBP deficiency impaired CSF1 signalling, and therefore macrophage differentiation and proliferation [42]. In light of this cross-talk and the key roles of TREM2+ macrophages in adipose and liver lipid metabolism [40, 41], we hypothesised that TREM2/TYROBP signalling may mediate some of the metabolic effects of CSF1-Fc treatment. TYROBP deficiency did compromise CSF1-induced differentiation in bone marrow-derived macrophages *in vitro* but there was no impact on any aspect of the CSF1-Fc response *in vivo*.

Liver and adipose macrophages are implicated in tissue adaptation to nutrient deprivation. Short-term fasting in mice led to CCR2-dependent infiltration of macrophages which engulfed excess lipids and regulated lipolysis to maintain tissue homeostasis [53]. We demonstrate that adipose macrophage expansion in response to CSF1 similarly regulates adipocyte size and adipose mass and show that the impacts of CSF1-Fc treatment and fasting were additive. In the liver, conditional deletion of the glucocorticoid receptor in macrophages reduced the induction of PPARγ target genes and ketogenesis in response to fasting [34]. This study did not include a control for potential toxic effects of expression of Cre recombinase in macrophages [54]. CSF1-Fc induces hepatic TNF expression [9, 10], which might be expected to impede the reported GR-driven ketogenesis. Here we found that neither expansion of tissue macrophages with CSF1-Fc, nor their depletion with anti-CSF1R, had any significant impact on any aspect of the hepatic response to fasting, despite a significant reduction in *Hmgcs* expression. Accordingly, we suggest that direct macrophage-hepatocyte interaction does not contribute to the regulation of ketogenesis. In this respect our conclusions are consistent with those of Merry *et al.* [55] who reported little effect of macrophage depletion using a CSF1R kinase inhibitor on glucose or lipid homeostasis.

## 5.0 Conclusions

CSF1-Fc treatment expands blood and tissue macrophages throughout the body and this effect is rapidly reversible. Treatment led to significant metabolic changes, notably liver growth, diversion of circulating glucose to the liver and the mobilisation of fat from adipose, which were also reversible. The response to CSF1-Fc treatment did not intersect with several known endocrine pathways regulating metabolism, and neither an increase, nor a decrease in tissue macrophages had any effect on normal hormonal regulation of metabolism in response to fasting. Despite rapid growth, CSF1-Fc-treated mice exhibited no change in appetence, and instead reduced activity and core body temperature, presumably to conserve energy and fuel growth. The mononuclear phagocyte system clearly functions as a whole-body system, with intrinsic homeostasis and system-wide communication mediated by CSF1R signals. Could cycles of CSF1 treatment be harnessed as a therapy alongside dietary modification to combat obesity-related pathology, mobilise fat and restore metabolic homeostasis? Combined with the established safety profile [56] and potential of this treatment to promote hepatic regeneration and tissue repair [9, 10] it is certainly worthy of consideration.

## Funding

This work was supported by: NHMRC Project Grant 1162171 and Ideas Grant 2012793 to KMI, NHMRC Project Grant APP1140064 and NHMRC fellowship APP1156489 to RGP, DARF MCR Fellowship to LGB, NHMRC Investigator Grant 2009750 to DAH, ARC Discovery Project DP20013245 to DAH and KMI, Australian Liver Foundation Pauline Hall Fellowship to SK.

## Declarations of interest

None.

## Author contributions

Conceptualisation: SK, DAH, KMI. Methodology: SK, MAS, BT, KAS, RP, KMS, DAH, KMI. Formal Analysis: SK, JM, LB, BT, KAS, RP, KMS, KMI. Investigation: SK, JM, LB, BT, KAS, OP, RP, KMS, KMI. Resources: HE, CS, LGB, ARP, DAH, KMI. Data Curation: SK, JM, KMI. Writing – Original Draft: DAH, KMI. Writing – Review and Editing: SK, DAH, KMI. Visualization: SK, BT, KAS, RGP, KMI. Supervision: ARP, DAH, KMI. Project Administration: KMI. Funding Acquisition: DAH, KMI.

## Acknowledgements

K.M.I., D.A.H., K.M.S. and A.R.P. are grateful for core laboratory support from the Mater Foundation. M.A.S. is supported by an Advance Queensland Industry Research Fellowship, Mater Foundation, Equity Trustees and the L G McCallam Est and George Weaber Trusts. We appreciate the support of the Preclinical Imaging, Biological Resources, Histology, Microscopy and Flow Cytometry Core Facilities at the Translational Research Institute. We are grateful to Dr David DeSouza, Metabolomics Australia, for performing metabolomic profiling and helpful advice and Mr Charles Ferguson for assistance with electron microscopy. The authors acknowledge the use of the Microscopy Australia Research Facility at the Centre for Microscopy and Microanalysis at The University of Queensland. We thank Q-Trace for kindly supplying the radiotracer for PET-CT.

## Notes

### Competing Interest Statement

The authors have declared no competing interest.

