## Supplementary Information for "Reversible expansion of tissue macrophages in response to macrophage colony-stimulating factor (CSF1) transforms systemic metabolism to fuel liver growth"

### Supplementary Methods.

#### Animal experiments.

Animal studies utilised 1 mg/kg P-CSF1-Fc[1] or 5 mg/kg Human CSF1-Mouse Fc conjugate (HM-CSF1-Fc, produced by Novartis, Switzerland), administered by sub-cutaneous injection, as indicated in results/figure legends. The two CSF1-Fc reagents were used interchangeably herein, based upon their availability. Previously reported minor differences in dose and precise time courses between the two CSF1-Fc reagents *in vivo* may be related to differential affinity for mouse CSF1R and/or impacts of the Fc region on circulating half-life [2]. The majority of studies were conducted in male mice on the C57BL/6J genetic background, unless otherwise indicated. C57BL6/J mice (6-10 weeks of age) were obtained from the Animal Resource Centre (ARC, Perth, Australia), or from our CSF1R-FusionRed breeding colony[3] (C57BL6/J background). Animals were housed in specific pathogen free facility on 12 hours dark cycle at 22-24°C with access to water and food ad libitum except during fasting. Fasting experiments were conducted from 8 am – 8 am (ZT1 in our animal facility). Anti-CSF1R mAb (AFS98, Cat#BE0213, Bioxcell, Assay Matrix, Australia) or control rat IgG (Cat#I4131, Sigma Aldrich, Australia) was administered by sub-cutaneous injection twice weekly for 4 weeks (400 µg/dose). For the fasting studies, fasting/feeding commenced/continued at 8 am on day 3 following the final dose and mice were sacrificed 24 h later. IGF1 (Bioxcell, Assay Matrix, Australia) or control saline was administered by intraperitoneal (IP) injection twice daily for 4 days (5 mg/kg/day). Mice were sacrificed on day 5. Beta blocker (propranolol hydrochloride Cat# P0884, Sigma Aldrich, Australia) or control saline was administered by IP injection from day -1 to 4 (10 mg/kg/dose). Mice were sacrificed on day 5. Anti-mouse IL6 (Clone MP5-20F3, Cat#BE0046, Lot#654520A2, BioxCell, Assay Matrix, Australia) or control rat IgG (Cat#I4131, Sigma Aldrich, Australia) was administered by IP injection on day -1 and 3 (0.5 mg/dose). Mice were sacrificed on day 5. Body composition was assessed using an NMR-based Minispec (Bruker LF50H). Blood glucose was measured from tail prick using a glucometer (SensoCard Australia). For the metabolic caging study, mice were surgically implanted with telemetry (VitalView/E-Mitter G, Starr LifeSciences, Oakmont, United States) in the peritoneal cavity and allowed to recover for 4 days before being housed in induvial metabolic cages (Phenomaster, TSE systems, Berlin, Germany). After 4 days acclimatisation, mice were injected with 5 mg/kg HM-CSF1-Fc or saline control for 4 days followed by 6 days recovery. Cage data (food and water intake, weight, oxygen and carbon dioxide) and telemetry data (temperature, activity) were recorded for the final 10 days of the experiment. At sacrifice, blood was collected by cardiac puncture for haematology analysis (Mindray BC-5000) and serum separation. For the high fat diet study, mice were maintained on normal chow and drinking water or HFD (based on American Institute of Nutrition (AIN)-93G, containing 23% fat, 0.19% cholesterol, Specialty Feeds SF15-059, Australia) and drinking water supplemented with 23 g/L fructose and 18.9 g/L sucrose (Sigma)[4]. Mice were fasted overnight (12 h) prior to sacrifice. TYROBP knock-out mice were generated by CRISPR/Cas9 gene editing. The fourth exon of the murine TYROBP gene was targeted using a mix of ribonucleic complexes formed by sgRNA (*in vitro* transcribed) and Cas9 enzyme (NEB #M0646) and a single-stranded DNA donor construct (Integrated DNA Technologies). The donor was designed to introduce silent mutations (enabling genotyping) at the target site, as well as a Tyr92Phe (TAT>TTT) mutation. Sequencing confirmed the presence of the silent mutations and the presence of a frameshift deletion at Tyr92 rendering the entire TYROBP signalling motif non-functional (LGB, manuscript in preparation). All animal experiments were approved by a University of Queensland Animal Ethics Committee.

**PET-CT Imaging and gamma counting.**

$^{18}$FDG was produced by Q-TRaCE, Department of Nuclear Medicine, Royal Brisbane and Women's Hospital, Australia. Mice were fasted for 6 h then injected with 4 MBq of [$^{18}$F]FDG diluted in 150 µl saline intravenously while under anaesthesia (2% isoflurane and oxygen mix). After 1 h of uptake, mice were anaesthetised and imaged inside a Molecubes β-Cube and Molecubes X-Cube (Molecubes, Belgium) for sequential PET and CT imaging. For PET, a 20 min static emission scan was performed, and images were reconstructed using an ordered-subset expectation maximization (OSEM) algorithm, with CT attenuation correction. For CT, parameters were 50 kV X-ray voltage, 75 µA current, 480 exposures at 85 ms each, continuous helical rotation, and images reconstructed at 200 µm isotropic voxel size using ISRA algorithm. PET-CT data was analysed and visualised using PMOD software v4.2 (PMOD Technologies, Switzerland). Standard uptake values (SUV) for specific volumes of interest were calculated. After imaging mice were euthanised by CO2 asphyxiation then tissues (liver, muscle, fat, brain, heart) were harvested and weighed for radioactivity measurement using a PerkinElmer 2480 Automatic Gamma Counter (Perkin Elmer, USA). The gamma counter was calibrated using samples of $^{18}$F of known concentration and measured activity was presented as percent injected dose per gram (%ID/g).

**Bone marrow derived macrophages.** Bone marrow obtained by flushing mouse femurs and tibias was seeded at 5e4 cells/well in a flat-bottomed 96 well tissue culture plate (Nunc) in complete RPMI culture medium containing 10 % FCS. Cells were treated with a dose response of recombinant human CSF1 (produced by the University of Queensland Protein Expression Facility) or recombinant mouse GM-CSF (RnD Systems), or without growth factor, for 7 days. On day 7 cell metabolism was quantified (as a surrogate of cell density and viability) by the addition of 0.025 mg/ml resazurin (Cayman Chemicals, Catalogue #14322) followed by incubation for a further 1 h. The fluorescent metabolite resorufin was quantified in a plate reader (Ex: 530-540 nm, Em: 585-595 nm).

**Histology.**

Whole mount imaging was performed on fresh tissue using a confocal microscope (FV3000, Olympus). Tissues for immunohistochemistry were fixed in 4% PFA for 24 hours and paraffin embedded. Epitope retrieval was performed in Diva Decloaker (Biocare Medical) followed by staining for Ki67 (Abcam, ab16667, lot GR3313195-28, 1:100) or F4/80 (Novus, NB600-404, Clone CI-A3-1, 1:400). Secondary detection was with DAKO Envision HRP reagents. Image quantification was performed from whole-slide digital images (VS200 scanner, Olympus) using ImageJ or Visiopharm software. Fresh frozen liver sections were sectioned at 7 µm using a Leica CM1950 cryostat followed by oil red O staining.

**Flow Cytometry**

Liver non-parenchymal cells were isolated as previously described[2]. Briefly, tissue disaggregation was performed by finely chopping liver samples (~1-2 g) in 10 ml digestion solution containing 1 mg/ml Collagenase IV (Thermofisher), 0.4 mg/ml dispase (Roche) and 20 µg/ml DNAse1 (Roche) and incubating at 37°C for 45 min on a rocking platform before mashing through a 70 µm filter (Falcon). The cell pellet was collected by centrifugation (400 g) and resuspended in an isotonic 30% Percoll solution to separate hepatocytes and non-parenchymal cells. Cells were stained for a panel of leukocyte markers [F4/80-AF647 (1:150), Cd11b-BV510 (1:200), Ly6G BV785 (1:200), (1:200), Tim4-PE-Cy7 (1:300), Ly6C-PE (1:300), CD3 APC-Cy7 (1:200), CD19 PE-Dazzle 594 (1:200) (Biolegend)] in buffer containing 2.4G2 supernatant to block Fc binding, washed and resuspended in buffer containing viability dye 7AAD (Life Technologies) for acquisition using a Cytoflex (Becton Dickinson). Live single cells were identified for phenotypic analysis by excluding doublets (FSC-A>FSC-H), 7AAD+ dead cells and debris. Single colour controls were used for compensation and unstained and fluorescence-minus-one controls were used to confirm gating. Data were analysed using FlowJo 10 (Tree Star). Cell counts were calculated by multiplying the frequency of the cell type of interest by the total mononuclear cell yield/g of disaggregated tissue.

**Gene expression analysis**

Tissue samples were collected in TRIzol (Sigma-Aldrich) for RNA extraction and cDNA synthesis (Bioline) according to manufacturer instructions. qPCR was performed using the SYBR Select Master Mix (Thermo Fisher Scientific) on an Applied Biosystems QuantStudio system. Primer pairs used in this study are as follows (5'-3'): Hprt F, GCAGTACAGCCCCAAAATGG, Hprt R, AACAAAGTCTGGCCTGTATCCAA;

Tbp F,CTCAGTTACAGGTGGCAGCA, Tbp R,ACCAACAATCACCAACAGCA;
Adgre1 F, CTGTCTGCTCAACCGTCAGGTA, Adgre1 R,
AGAAGTCTGGGAATGGGAGCTAA;
Ccl2 F, CAAGATGATCCCAATGAGTAGGC, Ccl2 R,
CTCTTGAGCTTGGTGACAAAAACTA;
Ccr2 F, GAACTTGAATCATCTGCAAAAACAAAT, Ccr2 R,
GGCAGGATCCAAGCTCCAAT;
Pdgfc F, AGCGCTGTGGAGGAAATTGT, Pdgfc R,
TTTTGGTCTCAACTGAAGGACCTC; Igf1 F, GCTGGTGGATGCTCTTCAGT, Igf1 R,
TCCGGAAGCAACACTCATCC; Ppara F, GGCTGCTATAATTTGCTGTGGAGAT ,
Ppara R,TTGCTCTGCAGGTGGAGCTT; Cyp4a10 F, CTTCTCAGGAGGAGCCAGGAA,
Cyp4a10 R, GTTCGAAGCGGAGCAGTGTC; Cyp4a14 F,
CCATTCTCAGGAGGATCAAGGAA , Cyp4a14 R, GTGGGATCTGGCAGCAATTC;
Hmgcs2 F, AAGGATGCTTCCCCAGGTT , Hmgcs2 R, TGCATCTCATCCACTCGTTC;
Socs2 F, GGTTGCCGGAGGAACAGTC , Socs2 R,
TCATACTTCCCCAGTACCATCCTG; Cish F, CCCTGTCCAGGCAGAGAATG , Cish R,
TAGAACCCCAGTACCACCCAGA; Il6 F, TCAGGAAATTTGCCTATTGAAA, Il6 R,
GGAAATTGGGGTAGGAAGGA; Apcs F, CATGGACAAGCTACTGCTTTGG, Apcs R,
CACAAATACTTTCCTCTTGAGGTCTG; Saa1/2 F, GGGCTGCTGAGAAAATCAGTG,

Saa1/2 R, CATGTCTGTTGGCTTCCTGGT; Trem2 F, ACCAACTTCAGATCCTCACTGGA,
Trem2 R, TGCAGGCTGGATTGACTCCT; TyrobpF, GTCAAGGGACAGCGGAAGG, TyrobpR, TTCTGGTCTCTGACCCTGAAGC.
Metrnl F, TGCCATCTGTACCAGTGACTTTGT, Metrnl R, CCGCAGGTAGATGACTGACACTT; Cd9 F, ACACCTACCAAAAGTTACGGAGCA, Cd9 R, AGCTATGCCACAGCAGTCCAAC; Arg1 F, GCCGATTCACCTGAGCTTTG, Arg1 R, TATCTAGTCCTGAAAGGAGCCCTGT; Clec10a F, TGGCCAACCTCAAGAACAACGG, Clec10a R, AGGCCACGACTTCTCAGACTCA; Retnl F, AGGAACTTCTTGCCAATCCAGCT, Retnl R, GCAGGAGGCCCATCTGTTCA; Mertk F, CCTGAGGACTGCTTGGATGAACTG, Mertk R, TCCAGCTGCAGCCTCAACAC; Timd4 F, ACTGTTTGCAGAGACACAAGAGG, Timd4 R, GAAGATCCCGTCTTCATCATCC.

**Metabolomics**

Metabolomics was performed by Metabolomics Australia (Melbourne). Metabolites were extracted from 50 µl serum using 150 µl methanol and 50 µl chloroform, including $^{13}$C-sorbitol and $^{13}$C,$^{15}$N-valine internal standards, and 30 µl of clarified sera was prepared for GC-MS analysis. Liver metabolites were extracted from 20-40 mg liver tissue (sampled from the same region of the left lateral lobe) in 600 µl methanol:MilliQ water (3:1), including $^{13}$C-sorbitol and $^{13}$C,$^{15}$N-valine internal standards. The tissue was cryomilled at 4°C. 110 µl chloroform was added to 440 µg homogenate which was transferred to a new tube and the sample thermomixed for 20 minutes then centrifuged at 13,000 rpm for 15 minutes. 30 µl supernatant was dried in inserts for GC-MS analysis. For both serum and liver pooled biological quality controls were compared by combining 50 µl aliquots of all samples. Quality controls clustered closely together and co-efficients of variation of randomly selected metabolites were within the acceptable range, demonstrating the robustness and reproducibility of the instrument and method. Samples were derivatised on-line using the AOC6000 autosampler robot in 25 µl methoxyamine hydrochloride in pyridine (30 mg/ml) 2hr at 37°C and 750 rpm agitation followed by the addition of 25 µl BSTFA with 1% TMCS and incubation for 1 h at 37°C and 750 rpm agitation, and 1 h at room temperature. Derivatised samples were analysed on Shimadzu TQ8050 using the Shimadzu Smart Metabolite database (comprising approximately 520 Multiple Reaction Monitoring profiles (MRMs) encompassing 350 metabolites. Data were processed using GC-MS Labsolutions Insight software.

**ELISA**

The serum concentration of triglyceride and β-Hydroxybutyrate were measured using colorimetric assay kits (Cat#10010303 and Cat#700190, respectively (Cayman Chemical)) according to the manufacturer’s instructions. The concentration of IL6 was measured using Mouse IL6 Quantikine ELISA Kit (M6000B, R&D Systems) and IGF1 level was measured using Mouse/Rat IGF-I/IGF-1 DuoSet ELISA (DY791, R&D Systems) according to manufacturer’s instructions. Serum insulin and glucagon levels were measured using Ultra-Sensitive Mouse Insulin ELISA Kit (Cat#90080, Crystal Chem) and Mercodia Glucagon ELISA kit (10-1271-01, Mercodia) respectively.

**Glycogen quantitation.**

Biochemical quantification of glycogen was performed as previously described[5].

### Data analysis

Data are presented as mean±s.d. Statistical tests were performed using GraphPad Prism 7.03. Data were analysed using t-tests, or 1 or 2-way ANOVAs with multiple testing correction. All tests were two-sided. $P<0.05$ was considered significant. Clustering analysis was performed in Graphia (www.graphia.app). Sample-sample clustering was performed at Pearson $r = 0.95$, using a Markov Cluster Algorithm (MCL) inflation of 1.3 for serum and liver. Metabolite-metabolite clustering was performed at Pearson $r = 0.75$, using a MCL inflation of 1.3.

### Data availability

Metabolomics data are available in the National Metabolomics Data Repository[6] under http://dx.doi.org/10.21228/M8TQ5G and http://dx.doi.org/10.21228/M8TQ5G for serum and liver, respectively.

**Supplementary Figures.**

**Supplementary Figure 1**

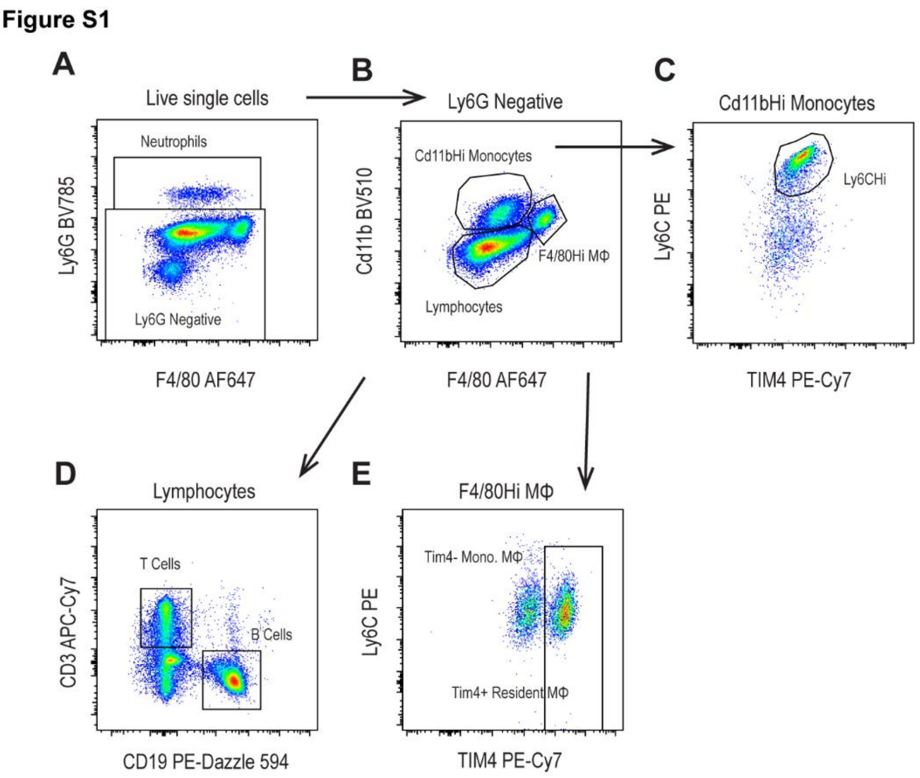


**Supplementary Figure 1. Liver leukocyte flow cytometry gating strategy.** Live, single cells were gated to identify Ly6G+ neutrophils (A). Ly6G negative cells were gated to identify lymphocytes, Cd11bHi/F4/80Int monocytes and F4/80Hi/Cd11bInt macrophages (MΦ) (B). Monocytes were separated into Ly6CHi and Ly6CLow subsets (C). Lymphocytes were gated to identify CD3+ T cells and CD19+ B cells (D). Macrophages were separated into Tim4+ and Tim4- subsets (E).

**Supplementary Figure 2**

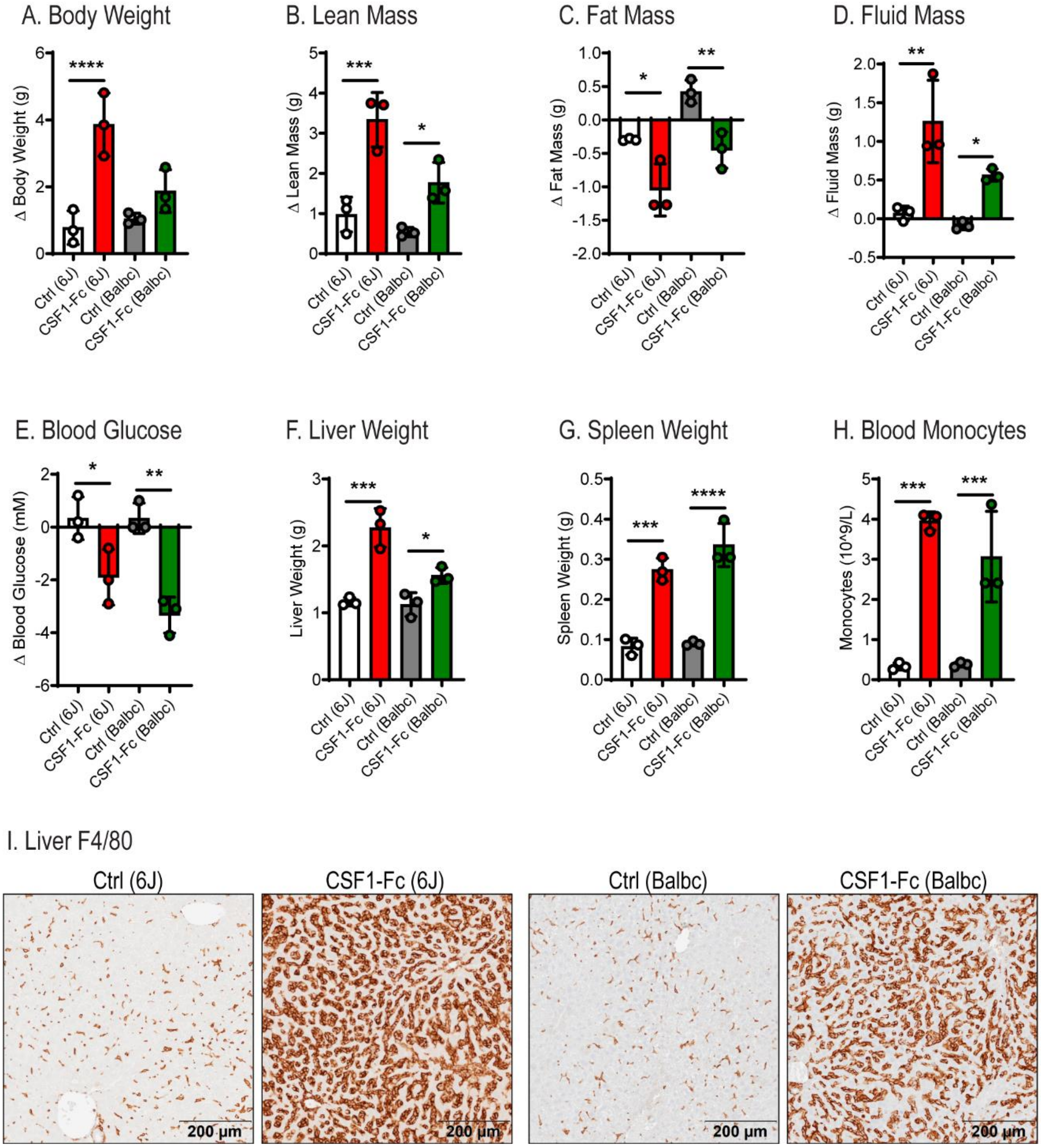


**Supplementary Figure 2. The impact of CSF1-Fc treatment is mouse strain independent.** Groups of 6 male C57BL6/J and BALB/C mice were treated with 5 mg/kg HM-CSF1-Fc or saline control daily for 4 days and sacrificed on day 5. Body weight (A), lean mass (B), fat mass (C), fluid mass (D), blood glucose (E), liver weight (F), spleen weight (G), blood monocyte count (H) at sacrifice. (I) Liver F4/80 staining. Data are presented as mean +/- s.d. are presented as mean +/- s.d. Ordinary one-way ANOVA with multiple comparisons.

**Supplementary Figure 3.**

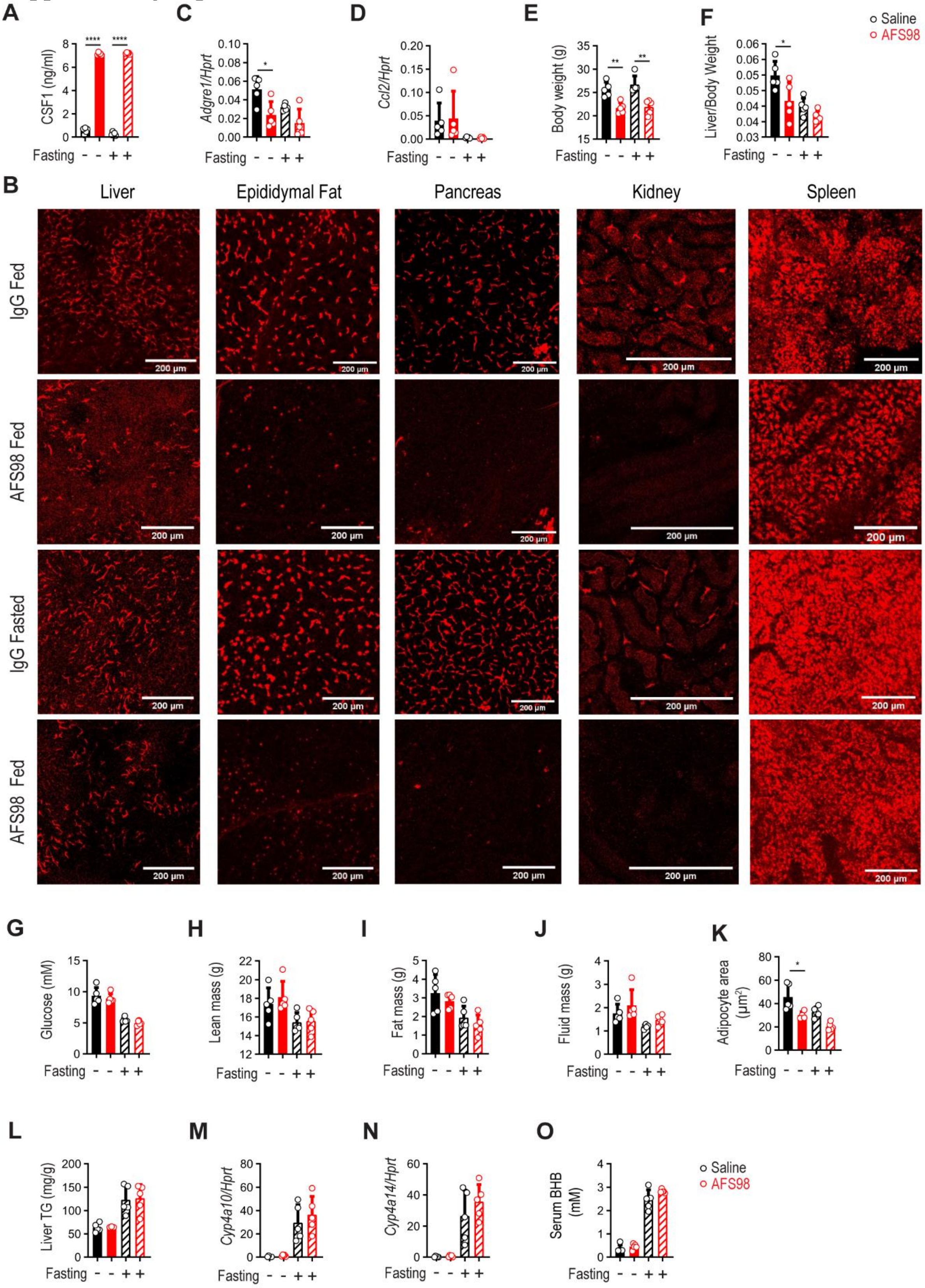


**Supplementary Figure 3. Systemic depletion of CSF1R-dependent macrophages does not impact the systemic response to fasting.** Groups of 5 male C57Bl/6J mice were treated twice weekly with 400 µg anti-CSF1R antibody (AFS98) or saline control by intraperitoneal injection for 2 weeks, followed by a 24 h fast initiated 3 days after the final dose prior to sacrifice. Circulating CSF1 (A) and whole-mount imaging of CSF1R+ macrophages (Csf1r-FusionRed reporter) in selected tissues (B) at sacrifice. Liver *Adgre1* (C) and *Ccl2* (D) mRNA expression, body weight (E) and liver weight (F) at sacrifice. Blood glucose (G), lean mass (H), fat mass (I), fluid mass (J), adipocyte area (K), liver triglycerides (TG) (L), liver *Cyp4a10* (M) and *Cyp4a14* (N) mRNA expression and serum β-hydroxybutyrate (BHB) (O) at sacrifice. Data are presented as mean +/- s.d. Two-way ANOVA with multiple comparisons (A-F, G-O).

**Supplementary Figure 4**

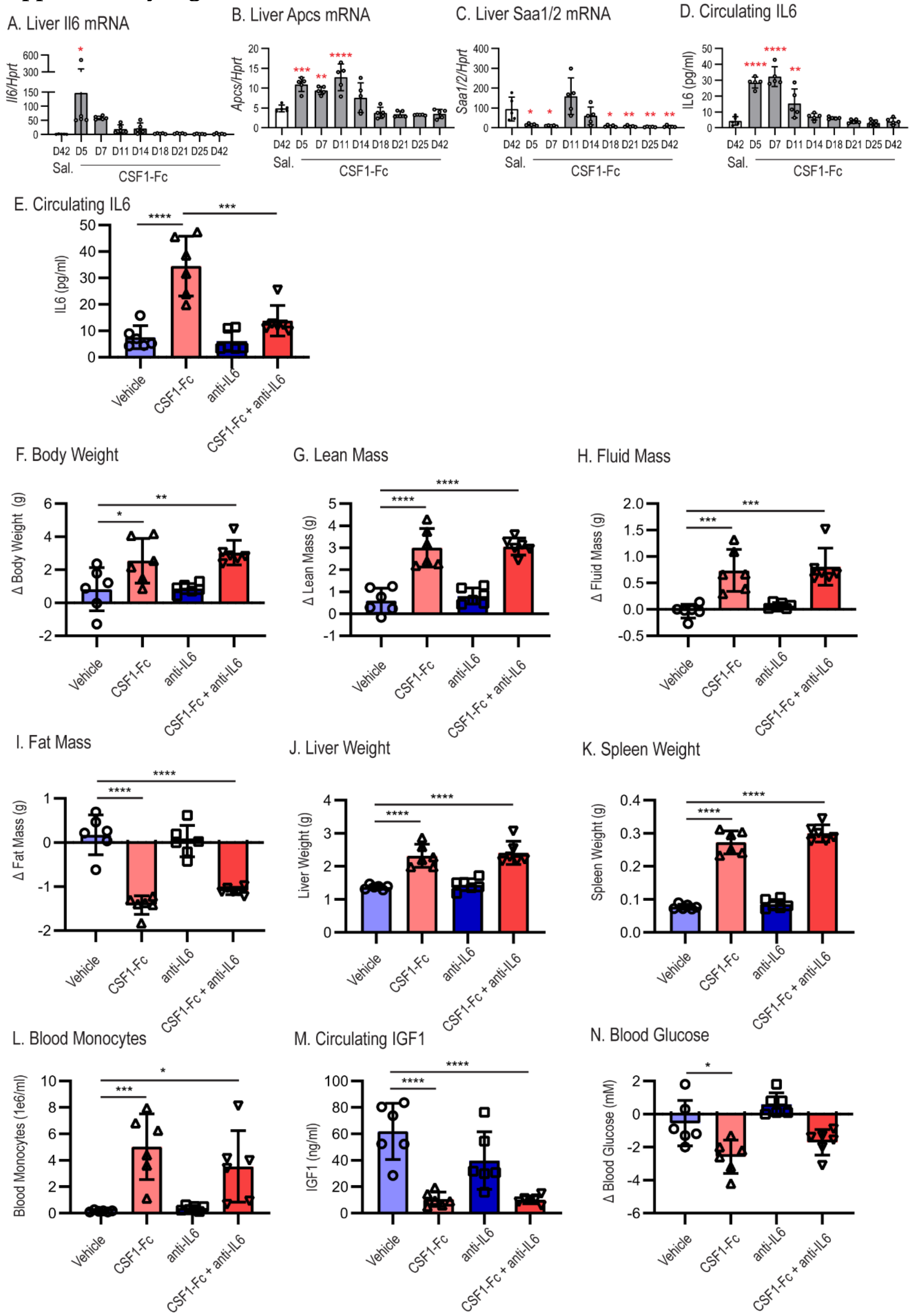

**Supplementary Figure 4. IL6 blockade does not impact the response to CSF1-Fc.** Groups of 6 male mice were administered 1 mg/kg P-CSF1-Fc for daily 4 days prior to sacrifice at the indicated timepoints up to 42 days. Control mice were treated with saline and sacrificed at 42 days. Liver *Il6* (A), *Apcs* (B), and *Saa1/2* (C) mRNA expression and serum IL6 (D) were measured at the indicated timepoints. Groups of male mice were treated with anti-IL6 antibody or control IgG (500 µg/kg by intraperitoneal injection) one day prior and on day 3 during an acute CSF1-Fc treatment regime. CSF1-Fc (5 mg/kg HM-CSF1-Fc) was administered daily for 4 days by sub-cutaneous injection and all groups were sacrificed on day 5. Serum IL6 (D), body weight (E), lean mass (F), fluid mass (G), fat mass (H), liver weight (I), spleen weight (J), blood monocyte count (K), circulating IGF1 (L) and blood glucose (M) at sacrifice. Data are presented as mean +/- s.d. Ordinary one-way ANOVA with multiple comparisons.

**Supplementary Figure 5**

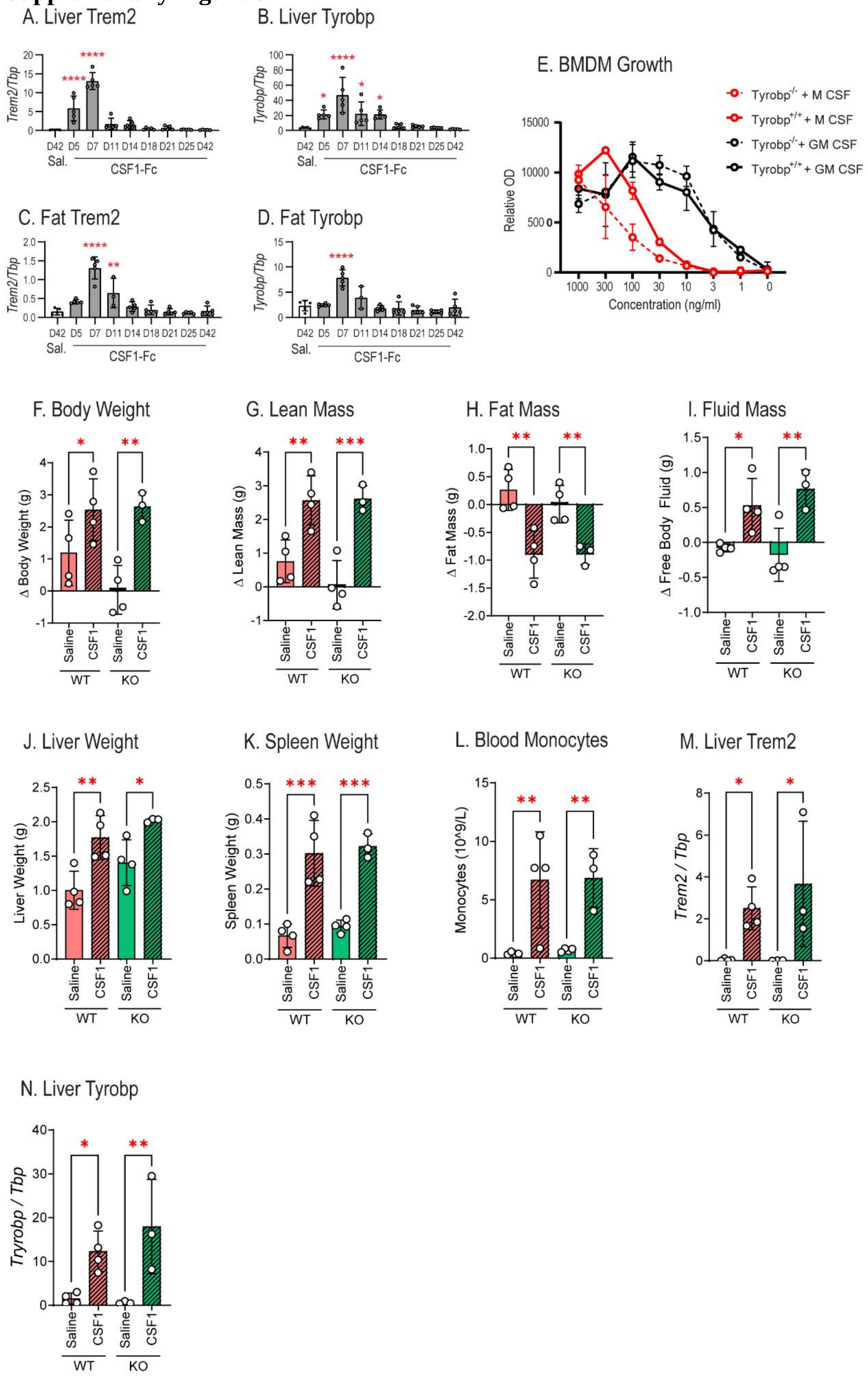

**Supplementary Figure 5. CSF1-Fc-induced fat mobilisation occurs in the absence of TYROBP signalling.** Groups of 5 male mice were administered 1 mg/kg P-CSF1-Fc for daily 4 days prior to sacrifice at the indicated timepoints up to 42 days. Control mice were treated with saline and sacrificed at 42 days. Liver *Trem2* (A) and *Tyrobp* (B) mRNA expression. Adipose *Trem2* (C) and *Tyrobp* (D) mRNA expression. (E) Bone marrow from WT and TYROBP KO mice was differentiated in CSF1 or GM-CSF at the indicated doses and cellular metabolic activity was assayed by resazurin reduction on day 7 (mean and s.d. of triplicates). (F-N) Groups of male WT and TYROBP KO mice were administered 5 mg/kg HM-CSF1-Fc or saline control daily for 4 days and sacrificed on day 5 (WT saline n=4, WT CSF1-Fc n=4) KO saline n=4, KO CSF1-Fc n=3). Body weight (F), lean mass (G), fat mass (H), fluid mass (I), liver weight (J), spleen weight (K), blood monocyte count (L), and liver *Trem2* (M) and *Tyrobp* (N) mRNA expression at sacrifice. Data are presented as mean +/- s.d. (A-D) Two-way ANOVA with multiple comparisons (F-N) Ordinary one-way ANOVA with multiple comparisons.

**Supplementary References.**
